# Awareness-dependent normalization framework of visual bottom-up attention

**DOI:** 10.1101/2021.04.18.440351

**Authors:** Shiyu Wang, Ling Huang, Qinglin Chen, Jingyi Wang, Siting Xu, Xilin Zhang

## Abstract

Although bottom-up attention can improve visual performance with and without awareness, whether they are governed by a common neural computation remains unclear. Using a modified Posner paradigm with backward masking, we found that both the attention-triggered cueing effect with and without awareness displayed a monotonic gradient profile (Gaussian-like). The scope of this profile, however, was significantly wider with than without awareness. Subsequently, for each subject, the stimulus size was manipulated as their respective mean scopes with and without awareness while stimulus contrast was varied in a spatial cueing task. By measuring the gain pattern of contrast-response functions, we observed changes in the cueing effect consonant with changes in contrast gain for bottom-up attention with awareness and response gain for bottom-up attention without awareness. Our findings indicate an awareness-dependent normalization framework of visual bottom-up attention, placing a necessary constraint, namely, awareness, on our understanding of the neural computations underlying visual attention.

## Introduction

Covert attention, the selective processing of visual information at a given location in the absence of eye movements, can be attracted automatically by an exogenous cue, known as visual bottom-up attention (*Corbetta and Shulman, 2002; Hegdé and Felleman, 2003; Kastner and Ungerleider, 2000; Koch and Ullman, 1985; Nakayama and Mackeben, 1989; Yantis and Jonides, 1984*). Numerous studies have demonstrated that this bottom-up attention-triggered attraction can improve visual performance with (*Beck and Kastner, 2005; Carrasco, 2011; Corbetta and Shulman, 2002; Kastner et al., 1997; Posner, 2016; Posner et al., 1980; Serences and Yantis, 2007*) and without awareness in various paradigms, such as visual backward masking (*Chen et al., 2016; Huang et al., 2020; Naccache et al., 2002; Zhang et al., 2012*), crowding (*Faivre and Kouider, 2011; Montaser-Kouhsari and Rajimehr, 2005*), and continuous flash suppression (*Bahrami et al., 2007; Hsieh et al., 2011; Jiang et al., 2006*), as well as sub-threshold presentation (*Bauer et al., 2009; Lin et al., 2009; Mulckhuyse and Theeuwes, 2010; Zhang and Fang, 2012*) and the patient with blindsight (*Kentridge et al., 1999a, 1999b, 2004*). However, it’s unclear whether there is a common neural computation governing bottom-up attention-triggered improvement in visual performance with and without awareness.

There has been a long-standing debate about the neural computations underlying visual bottom-up attention. Experiments examining how it modulates visual performance and neuronal activity in visual cortex have found disparate attentional effects on stimulus-evoked neural responses, such as the contrast-response function (CRF) that has often been used in the literature (*Martínez-Trujillo and Treue, 2002; Reynolds et al., 2000*). Some have reported that attentional selection primarily enhances neural responses to high-contrast stimuli (response gain, *Di Russo et al., 2001; Kim et al., 2007; Lee and Maunsell, 2009; Ling and Carrasco, 2006; McAdams and Maunsell, 1999; Morrone et al., 2002*), whereas others have reported that attentional selection primarily enhances neural responses to medium-contrast stimuli (contrast gain, *Li et al., 2008; Martínez-Trujillo and Treue, 2002; Reynolds and Chelazzi, 2004; Reynolds et al., 2000*). Still others have reported that attentional selection either enhances the entire contrast range or produces a combination of both response-gain and contrast-gain changes (*Buracas and Boynton, 2007; Huang and Dobkins, 2005; Murray, 2008; Pestilli et al., 2011; Williford and Maunsell, 2006*).

Crucially, these ostensibly conflicting results of the gain changes induced by visual attention can be explained by the normalization model of attention (*Boynton, 2009; Carandini and Heeger, 2012; Herrmann et al., 2010; Lee and Maunsell, 2009, 2010; Reynolds and Heeger, 2009; Reynolds et al., 1999*), which proposes that attention-triggered improvements on perception hinge on two critical factors: the stimulus size and the attention field size. Changes in the relative size of these two factors can tip the balance between neuronal excitatory and inhibitory processes, thereby resulting in response-gain changes, contrast-gain changes, or various combinations of the two (*Reynolds and Heeger, 2009*). Specifically, this model predicts that attention increases contrast gain when the stimulus is small and the attention field is large and increases response gain when the stimulus is large and the attention field is small. Remarkably, previous psychophysical (*Herrmann et al., 2010; Schallmo et al., 2020; Schwedhelm et al., 2016; Zhang et al., 2016*), electroencephalography (*Itthipuripat et al., 2014, 2019*), and voxel-based functional magnetic resonance imaging (*Hara et al., 2014*) studies have reported that the patterns of behavioral performance, steady-state visual evoked potentials, and voxel-averaged neurometric functions, respectively, are all consistent with the predictions of normalization model of attention. However, little is known regarding whether visual bottom-up attention with and without awareness is governed by this common neural computation: normalization and how awareness could modulate the gain changes induced by attentional selection.

Here, using a modified Posner paradigm with backward masking, we first manipulated the distance between the cue and probe (*Figure 1*) to measure the attention filed of visual bottom-up attention with and without awareness (*Distribution Experiments*). We found that the Posner cueing effect with and without awareness were both a monotonic gradient profile with a center maximum falling off gradually in the surround (Gaussian-like, *Figure 2*). Thus, for each subject, we fit their gradient profiles with a Gaussian function and used the FWHM (full width at half maximum) bandwidth of the Gaussian to quantify their scopes. The scopes of gradient profiles, however, were significantly wider with than without awareness, which offers a unique opportunity to change the size of the attentional scope relative to the stimulus size. Thus, for each subject, the stimulus size was manipulated as their respective mean scopes of visual bottom-up attention with and without awareness while stimulus contrast was varied in a spatial cueing task (*Normalization Experiments*). We measured the gain pattern of CRFs on the spatial cueing effect derived by visible or invisible cues and empirically revealed an interaction between awareness and visual bottom-up attention: gain modulation depended on awareness, with a change in the spatial cueing effect consonant with a change in contrast gain for visible cues and response gain for invisible cues. Additionally, using the classical normalization model of attention (*Reynolds and Heeger, 2009*), we successfully simulated the scopes of visual bottom-up attention with and without awareness. Our results thus support important predictions of the normalization model of visual bottom-up attention and further reveal its dependence on awareness.

**Figure 1.**
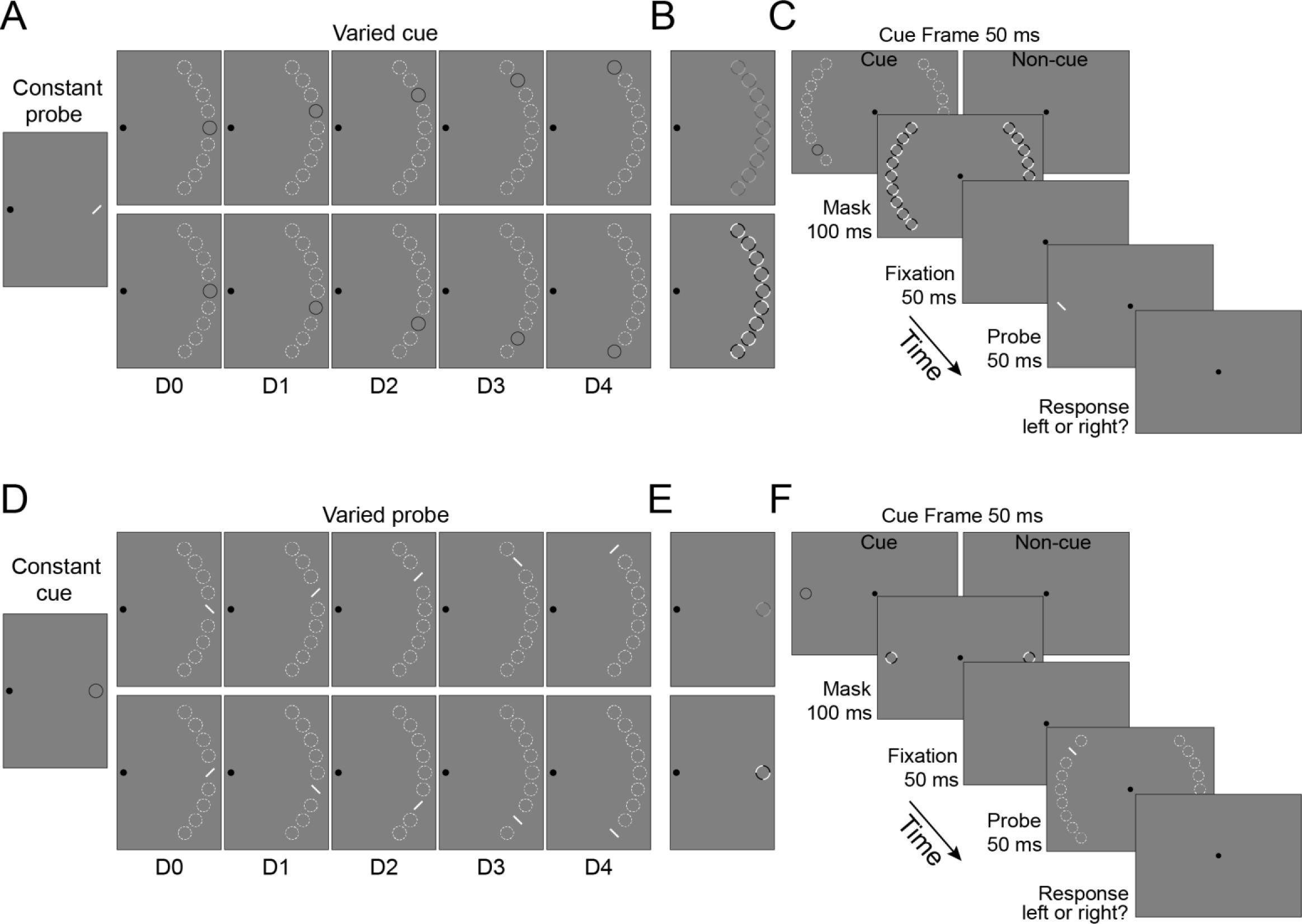
Stimuli and Psychophysical Protocol of Distribution Experiments. Each texture stimulus contained 18 positions, indicating by the dashed white circles (not displayed during the experiments), settled at an iso-eccentric distance from fixation (the black dot); a half of them were located in the left visual field and the other half was located in the right visual field. During Experiment 1 **(A)**, in each visual field (in this case, the right visual field), the probe, a tilted line (left), always appeared at the center of 9 positions while the exogenous cue, a low-luminance ring (right), appeared equiprobably and randomly at one of the 9 possible positions in the cue condition (83.33% of trials and 16.67% of trails for each distance) and was absent in the non-cue condition (16.67% of trials). The probe position was constant and the cue position varied, thus there were five possible distances between them, ranging from D0 (cue and probe at the same location) through D4 (cue and probe four items away from each other). During Experiment 2 **(D)**, in each visual field (in this case, the right visual field), the exogenous cue appeared (the cue condition, 50% of trials) or was absent (the non-cue condition, 50% of trials) in the center of 9 positions. The probe appeared equiprobably and randomly at one of the 9 possible positions. The cue position was constant and the probe position varied, there were also five possible distances between them (20% of trials for each distance), ranging from D0 through D4. Low- (top) and high- (bottom) contrast masks used for the visible and invisible conditions, respectively, in Experiments 1 **(B)** and 2 **(E)**. Psychophysical protocol of Experiments 1 **(C)** and 2 **(F)**. A cue frame with (the cue condition) or without (the non-cue condition) exogenous cue was presented for 50-ms, followed by a 100-ms mask (low- and high-contrast for visible and invisible conditions, respectively) and another 50-ms fixation interval. Then a probe line, orientating at 45° or 135° away from the vertical, was presented for 50-ms. Subjects were asked to press one of two buttons as rapidly and correctly as possible to indicate the orientation of the probe (45° or 135°).

**Figure 2.**
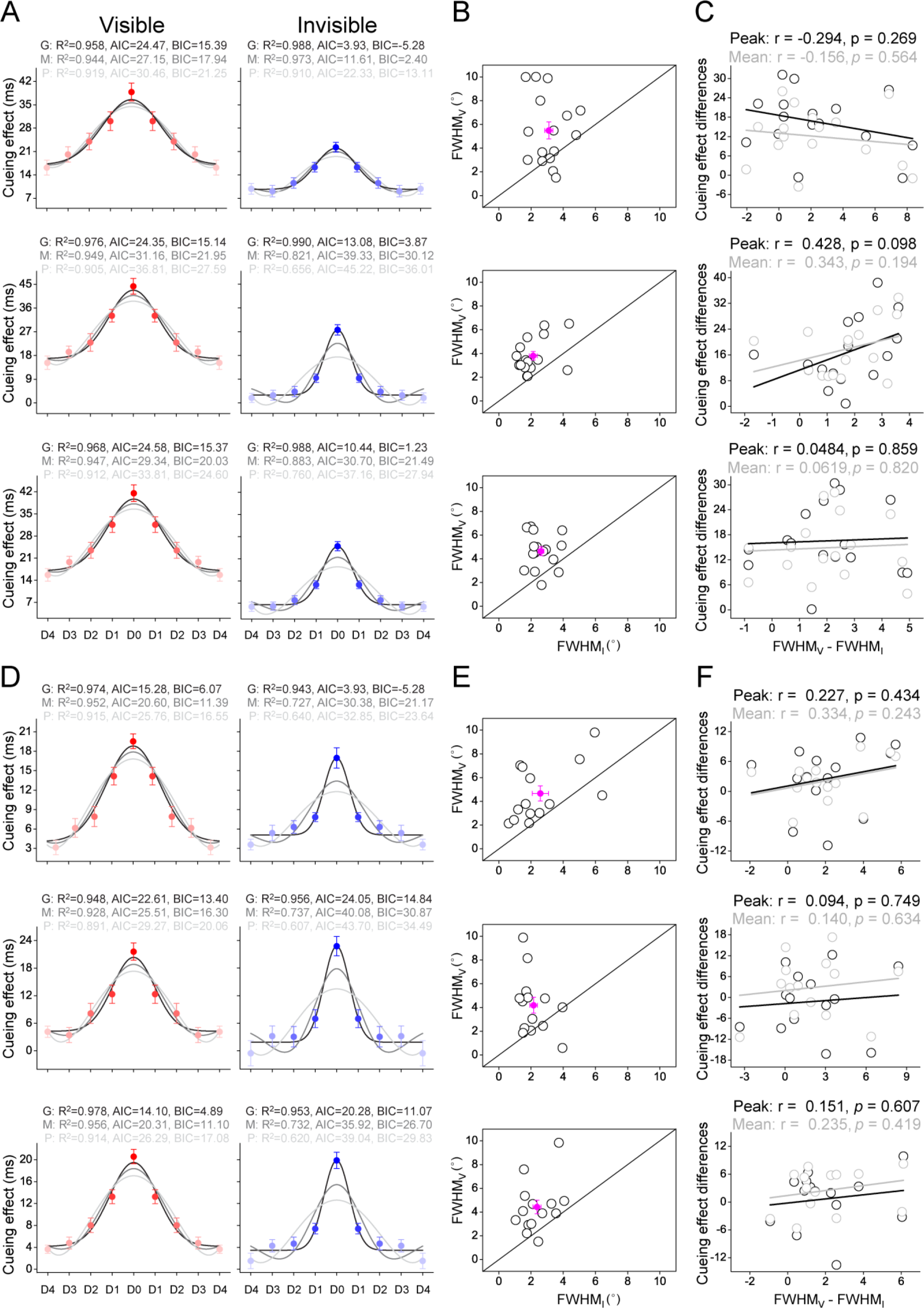
Results of Distribution Experiments. **(A)** and **(D)** The cueing effect of each distance (D0 to D4) in visible (left) and invisible (right) conditions for Experiment 1 (top), Experiment 2 (middle), and Experiments 1 & 2 (bottom), and the best fitting monotonic Gaussian function and two non-monotonic functions (Mexican Hat and Polynomial) to these cueing effects across distances. Each cueing effect was quantified as the difference between the reaction time of the probe task performance in the non-cue condition and that in the cue condition. Error bars denote 1 SEM calculated across subjects. G: Gaussian model; M: Mexican Hat model; P: Polynomial model; R^2^: R-squared; AIC: Akaike Information Criterion; BIC: Bayesian Information Criterion. **(B)** and **(E)** The fitted FWHM bandwidth of monotonic Gaussian model in visible and invisible conditions for Experiment 1 (top), Experiment 2 (middle), and Experiments 1 & 2 (bottom). FWHM_V_ and FWHM_I_: the fitted FWHM bandwidth of the Gaussian model in the visible and invisible conditions, respectively. Open symbols indicate individual subjects and a filled symbol indicate mean across subjects. Error bars denote 1 SEM calculated across subjects. **(C)** and **(F)** Correlations between the increased FWHM bandwidth and peak cueing effect (i.e., the cueing effect of D0, gray), and between the increased FWHM bandwidth and mean cueing effect (black) across distances, in the visible condition relative to the invisible condition, across individual subjects, in Experiment 1 (top), Experiment 2 (middle), and Experiments 1 & 2 (bottom).

## Results

### Distribution Experiments

The Distribution Experiment consisted of 3 experiments. In each visual field, the probe position was constant and the exogenous cue position varied in Experiment 1 (VCCP, *Figure 1A*), whereas Experiment 2 was a converse situation (CCVP, *Figure 1D*). In both Experiments 1 and 2, there were five possible distances between the exogenous cue and probe, ranging from D0 (cue and probe at the same location) through D4 (cue and probe four items away from each other). Subjects participated in Experiments 1 and 2 on two different days, and the order of the two experiments was counterbalanced across subjects. Experiment 3 checked the effectiveness of the awareness manipulation in both Experiments 1 and 2, and was always before them. Results showed that, our awareness manipulation was effective for both visible and invisible conditions (*Figure S1*). In both Experiments 1 (*Figure 1C*) and 2 (*Figure 1F*), each trial began with the fixation. A cue frame with (the cue condition) or without (the non-cue condition) exogenous cue was presented for 50-ms, followed by a 100-ms mask (low- and high-contrast masks rendered the exogenous cue visible or invisible to subjects, respectively) and another 50-ms fixation interval. Then a probe line, orientating at 45° or 135° away from the vertical, was presented for 50-ms. Subjects were asked to press one of two buttons as rapidly and correctly as possible to indicate the orientation of the probe (45° or 135°). There was no significant difference in the false alarm rate, miss rate, or removal rate (i.e., correct reaction times shorter than 200 ms and beyond three standard deviations from the mean reaction time in each condition were removed) across conditions (all *P* > 0.05, *Figure S2*). The cueing effect for each distance (D0 to D4) was quantified as the difference between the reaction time of the probe task performance in the non-cue condition and that in the cue condition.

*Figure 2A* shows the cueing effect of each condition for both Experiments 1 and 2; most of these cueing effects were significantly above zero, indicating that the bottom-up attention of the subject was attracted to the exogenous cue location, allowing them to perform more proficiently in the cue condition than the non-cue condition of the probe task. In both Experiments 1 and 2, a repeated measures ANOVA with awareness (visible and invisible) and distances (D0 to D4) as within-subjects factors showed that, the interaction between these two factors (Experiment 1: F_4, 60_ = 9.921, *p* < 0.001, 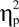 = 0.398; Experiment 2: F_4, 60_ = 3.36, *p* = 0.015, 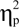 = 0.183), the main effect of awareness (Experiment 1: F_1, 15_ = 29.27, *p* < 0.001, 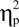 = 0.661; Experiment 2: F_1, 15_ = 72.26, *p* < 0.001, 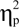 = 0.828), and the main effect of distances (Experiment 1: F_4, 60_ = 80.08, *p* < 0.001, 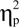 = 0.842; Experiment 2: F_4, 60_ = 86.30, *p* < 0.001, 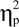 = 0.852) were all significant. Subsequent post hoc paired *t* tests revealed that the cueing effect decreased gradually with the distance in both Experiments 1 (the visible condition, D0 versus D1: t_15_ = 6.56, *p* < 0.001, 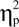 = 3.388, D1 versus D2: t_15_ = 3.68, *p* = 0.023, 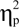 = 1.900, D2 versus D3: t_15_ = 4.36, *p* = 0.006, 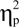 = 2.251, D3 versus D4: t_15_ = 3.18, *p* = 0.063, 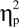 = 1.642; the invisible condition, D0 versus D1: t_15_ = 6.44, *p* < 0.001, 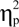 = 3.326, D1 versus D2: t_15_ = 4.13, *p* = 0.009, 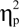 = 2.133, D2 versus D3: t_15_ = 1.60, *p* = 1.000, 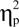 = 0.826, D3 versus D4: t_15_ = 0.60, *p* = 1.000, 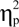 = 0.310) and 2 (the visible condition, D0 versus D1: t_15_ = 6.72, *p* < 0.001, 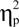 = 3.470, D1 versus D2: t_15_ = 5.70, *p* < 0.001, 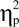 = 2.943, D2 versus D3: t_15_ = 1.28, *p* = 1, 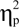 = 0.661, D3 versus D4: t_15_ = 2.08, *p* = 0.554, 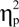 = 1.074; the invisible condition, D0 versus D1: t_15_ = 9.12, *p* < 0.001, 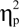 = 4.710, D1 versus D2: t_15_ = 2.27, *p* = 0.001, 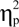 = 1.172, D2 versus D3: t_15_ = 0.50, *p* = 1.000, 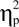 = 0.258, D3 versus D4: t_15_= 0.40, *p* =1.000, 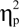 = 2.207). These results indicated that bottom-up attention-triggered cueing effects with and without awareness were both a monotonic gradient profile with a center maximum falling off gradually in the surround.

Subsequently, to further assess the shape of bottom-up attentional modulation, we fitted a monotonic model and two non-monotonic models to the average cueing effect across distances (D0 to D4) in both visible and invisible conditions. The monotonic model was implemented as the Gaussian function, and the two non-monotonic models were implemented as the Mexican Hat (i.e., a negative second derivative of a Gaussian function) and Polynomial functions (*Fang and Liu, 2019; Fang et al., 2019; Finke et al., 2008*). To compare these three models to our data, we first computed the Akaike information criterion (AIC, *Akaike, 1973*) and Bayesian information criterion (BIC, *Schwarz, 1978*) with the assumption of a normal error distribution. Then, we calculated the likelihood ratio (LR) and Bayes factor (BF) of the monotonic model (Gaussian) over non-monotonic models (Mexican Hat and Polynomial) based on AIC (*Burnham and Anderson, 2002*) and BIC (*Wagenmakers, 2007*) approximation, respectively. Results showed that, in both Experiments 1 and 2, the LR/BF (*Table 1, top*) strongly favored the Gaussian model over both the Mexican Hat and Polynomial models (*Figure 2A*). Notably, we also conducted similar model comparisons for each subject’ data and found that the Gaussian model was favored over both the Mexican Hat and Polynomial models in 11 and 10 for Experiment 1, in 9 and 9 for Experiment 2, out of 16 subjects, during the visible and invisible conditions, respectively (*Figure S4*). In addition, we pooled the data from Experiments 1 and 2 together and further provided the same qualitative conclusion. The LR/BF (*Table 1, top*) strongly favored the Gaussian model over both the Mexican Hat and Polynomial models (*Figure 2A, bottom*). The model comparison based on fitting individual data also demonstrated that the Gaussian model was favored over both the Mexican Hat and Polynomial models in 12 and 10 out of 16 subjects during the visible and invisible conditions, respectively (*Figure S4*). These results further constituted strong evidence for the monotonic gradient profile of visual bottom-up attention with and without awareness.

**Table 1.**
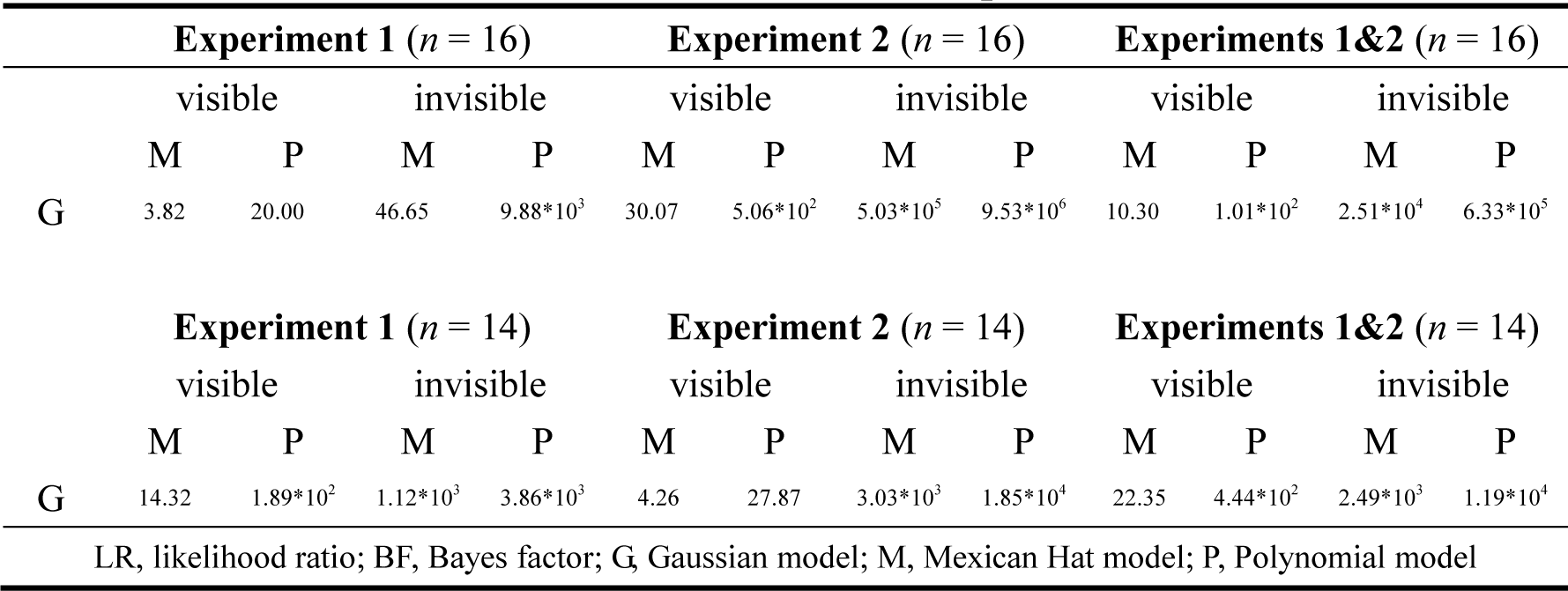
LR/BF of model comparisons.

Our results indicated that the spatial focus of visual bottom-up attention with and without awareness were best explained by the monotonic (Gaussian) rather than the non-monotonic models (Mexican Hat and Polynomial). To quantitatively examine the scope of bottom-up attentional modulation, we fitted the cueing effects from D0 to D4 with a Gaussian function and used the FWHM bandwidth of the Gaussian to quantify their scopes. Results showed that the fitted FWHM bandwidth was significantly larger in the visible than the invisible condition for both Experiments 1 (t_15_ = 3.015, *p* = 0.009, 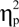 = 1.557, *Figure 2B, top*) and 2 (t_15_ = 4.863, *p* < 0.001, 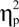 = 2.511, *Figure 2B, middle*), as well as for the pooled data from two experiments (t_15_ = 4.745, *p* < 0.001, 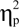 = 2.450, *Figure 2B, bottom*), indicating a wider scope of bottom-up attention with than without awareness. Notably, this awareness-dependent scope of bottom-up attention here could be explained by the difference in cueing effect between the visible and inviable conditions. To examine this issue, we calculated the correlation coefficients between our fitted FWHM bandwidths and cueing effects across individual subjects. If a wider scope of visual bottom-up attention with than without awareness is derived by a greater cueing effect in the visible than the invisible condition, then we would observe a significant correlation between these two measures across individual subjects. However, for Experiments 1 and 2, as well as for the pooled data from the two, compared to the invisible condition, the increased FWHM bandwidth was not significantly correlated with the increased peak cueing effect (i.e., the cueing effect of D0, Experiment 1: r = -0.294, *p* = 0.269, 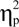 = 0.086; Experiment 2: r = 0.428, *p* = 0.098, 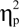 = 0.183; Experiments 1 & 2: r = 0.0484, *p* = 0.859, 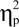 = 0.002) or the mean of cueing effects across distances (Experiment 1: r = -0.156, *p* = 0.564, 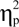 = 0.024; Experiment 2: r = 0.343, *p* = 0.194, 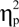 = 0.118; Experiments 1 & 2: r = 0.0619, *p* = 0.820, 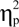 = 0.004) in the visible condition (*Figure 2C*), which against the cueing effect explanation.

More importantly, to directly exclude this explanation, we examined the scope of bottom-up attention during visible and invisible conditions with no significant difference in the cueing effect between the two conditions. We manipulated the cueing effect of visible condition by decreasing the luminance of its cue (which was still visible to subjects, *Figure S1*). Fourteen of our 16 subjects repeated Distribution Experiments using these low luminance cues and the same repeated measures ANOVA indicated that our manipulation was effective by showing that, in both Experiments 1 and 2, the main effect of awareness (Experiment 1: F_1, 13_ = 3.283, *p* = 0.093, 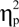 = 0.202; Experiment 2: F_1, 13_ = 1.472, *p* = 0.247, 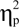 = 0.102) was not significant (*Figure 2D*). Our model comparisons also provided the same qualitative conclusion that the Gaussian model was strongly favored over both the Mexican Hat and Polynomial models with the LR/BF (*Table 1, bottom*), based on both the group (*Figure 2D*) and individual (*Figure S4*) data. More importantly, we confirmed that the fitted FWHM bandwidth was significantly larger in the visible than the invisible condition for both Experiments 1 (t_13_ = 3.732, *p* = 0.003, 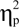 = 2.070, *Figure 2E, top*) and 2 (t_13_ = 2.561, *p* = 0.024, 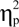 = 1.421, *Figure 2E, middle*), as well as for the pooled data from the two (t_13_ = 3.752, *p* = 0.002, 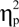 = 2.081, *Figure 2E, bottom*). Additionally, these increased FWHM bandwidths in the visible condition relative to the invisible condition weren’t significantly predicted by the increased peak cueing effect (Experiment 1: r = 0.227, *p* = 0.434, 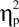 = 0.052; Experiment 2: r = 0.094, *p* = 0.749, 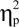 = 0.009; Experiments 1 & 2: r = 0.151, *p* = 0.607, 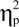 = 0.023) or the mean of cueing effects across distances (Experiment 1: r = 0.334, *p* = 0.243, 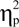 = 0.112; Experiment 2: r = 0.140, *p* = 0.634, 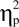 = 0.020; Experiments 1 & 2: r = 0.235, *p* = 0.419, 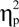= 0.055) (*Figure 2F*). Thus, our findings indicate a gradient profile of visual bottom-up attention with and without awareness, and show a wider scope of visual bottom-up attention with than without awareness.

### Normalization Experiments

Our Distribution Experiments demonstrated an awareness-dependent scope of visual bottom-up attention, which offers a unique opportunity to change the size of the attention field relative to the stimulus, differentially modulating the gain of bottom-up attentional selection. Thus, for each subject, the diameter of grating (*Figure 3A*) was manipulated as their respective mean FWHM bandwidth of the Gaussian with and without awareness, i.e., the diameter of grating = (FWHM_V_ + FWHM_I_) / 2, where FWHM_V_ and FWHM_I_ are the fitted FWHM bandwidth of the Gaussian model for the visible and invisible conditions in Distribution Experiments, respectively. Under this configuration, the attentional field was larger and smaller than the stimulus size for the visible and invisible cues, yielding a pattern that qualitatively resembled contrast gain or response gain, respectively (*Figure 3B*). To examine these predictions, we used a modified version of the Posner paradigm to measure the cueing effect induced by the visible or invisible cue, as shown in *Figure 3C*. In both two conditions, an exogenous cue, a low-luminance ring, randomly appeared at the center of 9 positions in left or right hemifield with equal probability, followed by a 100-ms mask (low- and high-contrast for visible and invisible conditions, respectively) and another 50-ms fixation interval. Then, a pair of gratings was presented for 33 ms in the left and right hemifields and subjects were asked to press one of two buttons to indicate the orientation of one of two gratings; each was presented at five different contrasts (0.02, 0.08, 0.15, 0.40, and 0.70, the contrasts of both gratings were identical on any given trial and covaried across trials in random order). A response cue at gratings offset indicated the target grating, yielding congruent cue (the exogenous cue matched the response cue, half the trials) and incongruent cue (mismatched, half the trials) conditions (*Figure 3C*). Comparing performance accuracy (*d’*) for congruent and incongruent trials revealed the spatial cueing effect for each target contrast.

**Figure 3.**
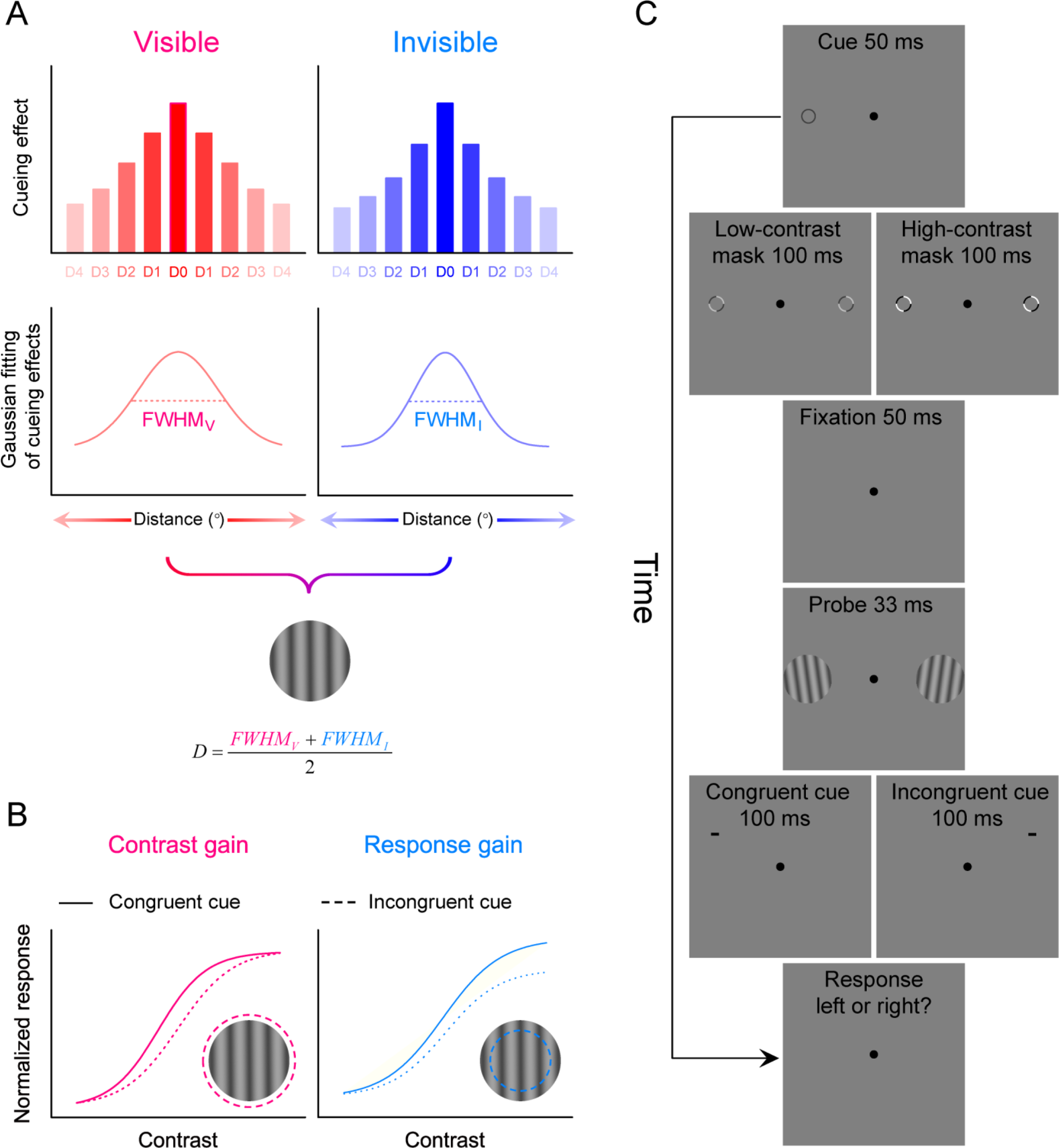
The Normalization Model of Attention Modulated by Awareness and Psychophysical Protocol in Normalization Experiments. **(A)** An example of grating probes manipulated from Distribution Experiments. For each subject, the diameter of grating = (*FWHM_V +_ FWHM_I_*) / 2, where *FWHM_V_* and *FWHM_I_* are their fitted FWHM bandwidth of the Gaussian model with and without awareness during Distribution Experiments (Experiments 1 & 2), respectively. **(B)** Relative to the stimulus size, the attention field was broadened by visible exogenous cues (left). Under this configuration, the normalization model predicts a contrast-gain shift, with the largest effects occurring at mid-contrasts and little to no effect at low and high contrasts. Relative to the stimulus size, the attention field was narrowed by invisible cues (right). Under this configuration, the normalization model predicts a response-gain shift, with the largest effects occurring at high contrasts and little to no effect at low and mid-contrasts. The dashed circles indicate simulated attention field size. **(C)** Psychophysical protocol. The exogenous cue randomly appeared at the center of 9 positions in left or right hemifield with equal probability, followed by a 100-ms mask (low- and high-contrast for visible and invisible conditions, respectively) and another 50-ms fixation interval. Then, a pair of gratings (with identical contrasts) was presented for 33 ms in the left and right hemifields, one of which was the target. Subjects were asked to press one of two buttons to indicate the orientation of the target grating (leftward or rightward tilted) and received auditory feedback if their response was incorrect. The target grating was indicated by a peripheral 100 ms response cue above one of the grating locations, but not at the grating location to avoid masking. A congruent cue was defined as a match between the exogenous cue location and response cue location (half the trials); an incongruent cue was defined as a mismatch (half the trials).

The mean *d’* plotted as psychometric functions of stimulus contrast and awareness (visible and invisible) are shown in *Figure 4A*: the visible condition yielded a pattern that qualitatively resembled contrast gain, and the invisible condition yielded a pattern that qualitatively resembled response gain (see also *Figure S5A*). The measured psychometric function for awareness (visible and invisible) and trial conditions (congruent and incongruent) was fit with the standard Naka–Rushton equation (*Naka and Rushton, 1966*). The two parameters *c_50_* (the contrast yielding half-maximum performance) and *d’*_max_ (asymptotic performance at high-contrast levels) determined contrast gain and response gain, respectively. The exponent *n* (slope) was fixed at 2 in the current analysis (*Carandini and Heeger, 2012; Herrmann et al., 2010; Reynolds and Heeger, 2009; Zhang et al., 2016*). The *d’* _max_ for awareness (visible and invisible) and trial conditions (congruent and incongruent) are shown in *Figure 4* and were submitted to a repeated-measures ANOVA with awareness and trial condition as within-subjects factors. The main effect of awareness (F_1, 12_ = 8.915, *p* = 0.011, 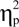 = 0.426), the main effect of the trial condition (F1, 12 = 70.366, *p* < 0.001, 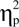 = 0.854) and the interaction between these two factors (F_1, 12_ = 71.311, *p* < 0.001, 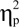 =0.856) were all significant. Post hoc paired *t* tests showed that *d’* _max_ of congruent trials was higher than that of incongruent trials for the invisible condition (t_12_ =12.166, *p* < 0.001, 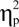 = 7.024, *Figure 4C, left*), but not for the visible condition (t_12_ =1.784, *p* = 1.000, 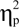= 1.030, *Figure 4B, left*); *d’*max for the invisible condition was higher than that for the visible condition in the congruent trials (t_12_ = 5.163, *p* < 0.001, 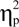 = 2.981), but not in the incongruent trials (t_12_ = 1.098, *p* = 0.294, 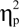 = 0.634). Similarly, for the *c*_50,_ the main effect of awareness (F1, 12 = 5.468, *p* = 0.037, 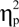 = 0.313), the main effect of trial condition (F1, 12 = 45.342, *p* < 0.001, 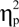 = 0.791) and the interaction between these two factors (F_1, 12_ = 52.415, *p* < 0.001, 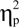 = 0.814) were all significant. Post hoc paired *t* tests showed that *c*_50_ of congruent trials was lower than that of incongruent trials for the visible condition (t_12_ = -9.303, *p* < 0.001, 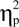 = -5.371, *Figure 4D, left*), but not for the invisible condition (t_12_ = -0.577, *p* = 0.575, 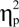 = -0.333, *Figure 4E, left*); *c*_50_ for the visible condition was lower than that for the invisible condition in the congruent trials (t_12_ = -5.074, *p* < 0.001, 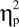 = -2.929), but not in the incongruent trials (t_12_ = 0.023, *p* = 0.982, 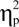 = 0.013). These results thus suggest that gain modulation of bottom-up attentional selection depends on awareness.

**Figure 4.**
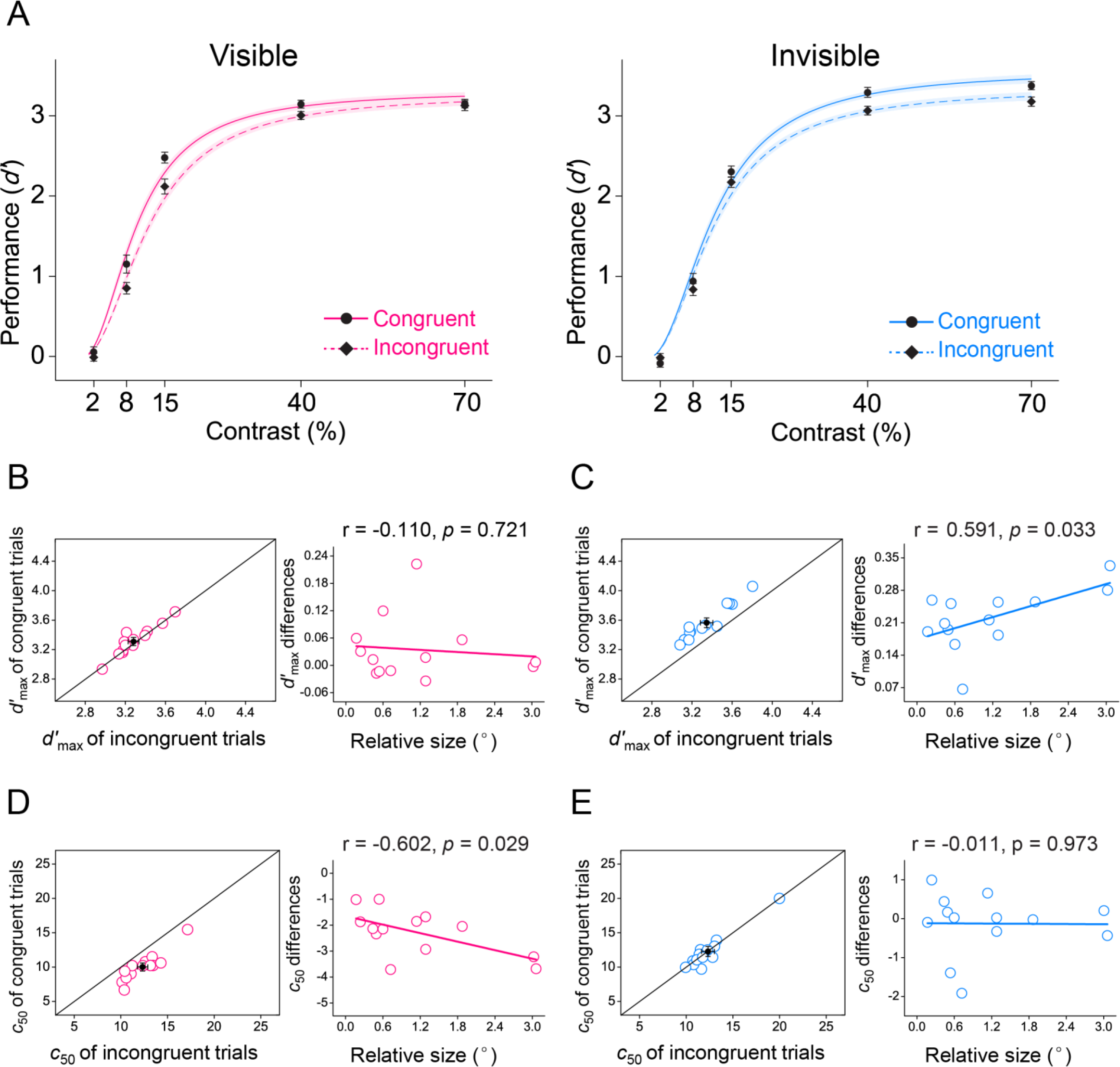
Effects of Visible and Invisible Cues on Performance (d’) as a Function of Contrast. **(A)** Mean *d’* plotted as psychometric functions of stimulus contrast and awareness (visible, left; invisible, right) for congruent and incongruent trials. Error bars denote 1 SEM calculated across subjects. *d’*_max_ for congruent and incongruent trials in the visible **(B)** and invisible **(C)** conditions, and correlations between the relative size of attention field to the stimulus [i.e., (FWHM_V_ - FWHM_I_) / 2, where FWHM_V_ and FWHM_I_ are the fitted FWHM bandwidth of the Gaussian model for the visible and invisible conditions, respectively] and the *d’*_max_ differences between congruent and incongruent trials across individual subjects. Open symbols indicate individual subjects and a filled symbol indicate mean across subjects. Error bars denote 1 SEM calculated across subjects. *d’*_max_: asymptotic performance at high contrast levels. *c*_50_ for congruent and incongruent trials in the visible **(D)** and invisible **(E)** conditions, and correlations between the relative size of attention field to the stimulus and the *c*_50_ differences between congruent and incongruent trials across individual subjects. Open symbols indicate individual subjects and a filled symbol indicate mean across subjects. Error bars denote 1 SEM calculated across subjects. *c*_50_: the contrast yielding half maximum performance.

To evaluate further the role of awareness in the gain modulation of visual bottom-up attention, we calculated the correlation coefficients between the relative size of the attention field to the stimulus [i.e., (FWHM_V_ - FWHM_I_) / 2, where FWHM_V_ and FWHM_I_ are the fitted FWHM bandwidth of the Gaussian model for the visible and invisible conditions, respectively] and psychophysical measures (*d’* _max_ and *c*_50_) across individual subjects. The relative size of the attention field to the stimulus in the visible condition significantly correlated with the *c*_50_ difference between congruent and incongruent trials (r = -0.602, *p* = 0.029, 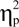= 0.362, *Figure 4D, right*), but not with the *d’* _max_ difference between congruent and incongruent trials (r = -0.110, *p* = 0.721, 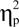 = 0.012, *Figure 4B, right*). Conversely, the relative size of the attention field to the stimulus in the invisible condition significantly correlated with the *d’* _max_ difference between congruent and incongruent trials (r = 0.591, *p* = 0.033, 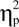 = 0.349, *Figure 4C, right*), but not with the *c*_50_ difference between congruent and incongruent trials (r = -0.011, *p* = 0.973, 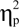= 0.0001, *Figure 4E, right*). These results thus demonstrate a close relationship between awareness and gain modulation of visual bottom-up attentional selection (response gain and contrast gain changes in psychophysical performance).

In addition, to further confirm this awareness-dependent normalization framework of visual bottom-up attention, we simulated our empirical data with the normalization model of attention (*Figure 5A* and *Figure S5B*) using custom Matlab scripts based on the code of Reynolds and Heeger (*2009*) with 4 free parameters: the gain of attention [*A(x,θ)*], separately optimized for visible and invisible conditions, the normalization constant *σ*, and a scaling parameter to linearly scale simulated values to performance (*d’*). Given the simulated attention fields [*A*(*x*,*θ*)] are in arbitrary units; only the relative values are meaningful (*Reynolds and Heeger, 2009*), in both the visible and invisible conditions, we thus calculated the correlation coefficients between the simulated and experimental attention fields (i.e., the FWHM) across individual subjects. In both two conditions, the simulated attention fields (marginally) significantly correlated with the experimental attention fields (the visible conditions: r = 0.792, *p* = 0.001, 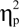 = 0.627; the invisible conditions: r = 0.505, *p* = 0.079, 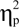 = 0.255, *Figure 5B*), further conforming that manipulating subjects’ awareness could modulate the field of visual bottom-up attention, which, in turn, affected its normalization processes.

**Figure 5.**
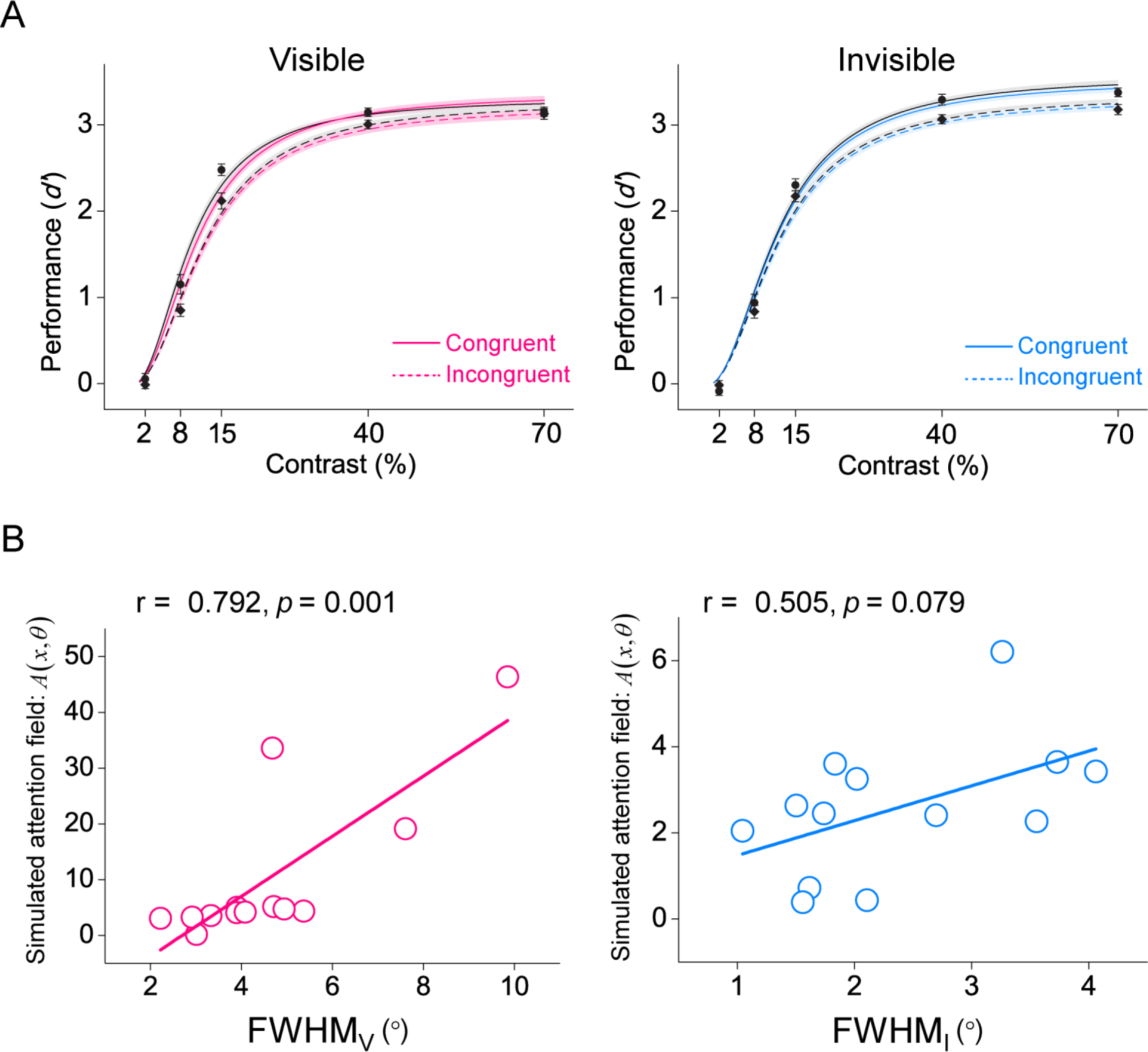
Model Predictions of Contrast Response Functions for the Normalization Model of Attention. **(A)** Data points with plus/minus one standard error are the mean performance (*d’*) across subjects for awareness (visible and invisible), cue validity (congruent and incongruent), and stimulus contrasts. The fitted performances (*d’*) using the standard Naka–Rushton equation and the original normalization model of attention with 4 free parameters are indicated by the black and colored lines, respectively. **(B)** Correlations between the simulated attention field [*A*(*x*,*θ*)] of the original normalization model of attention with 4 free parameters and the experimental attention field in the visible (left) and invisible (right) conditions, across individual subjects. Note that the simulated attention fields [*A*(*x*,*θ*)] are in arbitrary units; only the relative values are meaningful.

## Discussion

We examined the underlying neural computations of visual bottom-up attention with and without awareness and found the following results. First, we found support for previous neurophysiological (*Connor et al., 1996, 1997; Schall and Hanes, 1993*), psychophysical (*Downing 1988; Handy et al., 1996; Henderson and Macquistan, 1993; LaBerge, 1983; Mangun and Hillyard, 1988; Posner, 2016; Robertson et al., 2013; Shulman et al. 1985*), eye movement (*Tkacz-Domb and Yeshurun, 2018*), electroencephalographic (*Couperus and Lydic, 2019; Eimer, 1997; Mangun and Hillyard, 1988*), and functional magnetic resonance imaging (*Brefczynski-Lewis et al., 2009*) studies, indicating that the bottom-up attention-triggered improvements in visual performance with and without awareness were both a monotonic gradient profile with a center maximum falling off gradually in the surround (Gaussian-like). Second, however, the scope of this gradient profile was significantly wider with than without awareness, which offers a unique opportunity to change the size of the bottom-up attentional scope relative to the stimulus size. Thus, for each subject, the stimulus size was manipulated as their respective mean scopes of bottom-up attention with and without awareness while stimulus contrast was varied. By measuring the gain pattern of CRFs on the spatial cueing effect derived from visible or invisible exogenous cues, we observed a change in the spatial cueing effect consistent with a change in contrast gain for visible cues and in response gain for invisible cues. Finally, using the classical normalization model of attention (*Reynolds and Heeger, 2009*), we successfully simulated the scopes of visual bottom-up attention with and without awareness, indicating an awareness-dependent normalization framework of visual bottom-up attention. In addition, our results cannot be explained by the eye movement since subjects’ eye movements were small and their eye position distributions were statistically indistinguishable for visible and invisible conditions (*Figure S3*).

### Gradient profile of visual bottom-up attention with and without awareness

Compared to a monotonic gradient profile evident in previous and our studies, a number of neurophysiological (*Moran and Desimone, 1985; Schall and Hanes, 1993; Schall et al., 2004*), psychophysical (*Bahcall and Kowler, 1999; Mounts, 2000; Müller et al., 2005*), and brain imaging (*Boehler et al., 2009, 2011; Hopf et al., 2006, 2010; Müller and Kleinschmidt, 2004*) studies, as well as a computational model (*Tsotsos et al., 1995, 2008*) have reported a center–surround (i.e., the Mexican Hat) profile where a zone of sensory attenuation surrounds a center region of facilitation. We suggested that this striking discrepancy in the literature findings might be due to two different factors. One is the experimental task or paradigm. Specifically, for example, in Hopf et al. (*2006*) where inhibition was seen surrounding the locus of spatial attention, subjects were asked to search for a target (i.e., the exogenous cue in our study), which appeared randomly to change the spatial focus of attention and thus was task-relevant. While they measured the event-related magnetic field response elicited by a task-irrelevant probe that appeared or was absent with equal probability. In our study, by contrast, we used a modified Posner paradigm to measure the probe’s attentional effect induced by the exogenous cue. Subjects were asked to discriminate the orientation (45° or 135°) of the probe (*Figure 1*), thus the exogenous cue was task-irrelevant while the probe was task-relevant. Obviously, the probe was the distractor (task-irrelevant) and the target (task-relevant) in Hopf et al.’s and our studies, respectively. Directing attention to the target could conflate perceptual and post-perceptual mechanisms of attention, thus eliminating or strongly attenuating the suppression effect.

The other that may influence the spatial profile of visual bottom-up attention is whether the exogenous cue was presented with (e.g., Hopf et al., *2006*) or without (e.g., the current study) the distractors. As is known to all, the biased competition model of attention (*Desimone and Duncan, 1995*) proposes that attention operates when multiple stimuli compete for access to neural representation, and this competition occurs when multiple stimuli fall within a neuron’s receptive field. In this case, distractors within a receptive field are suppressed while attended stimuli are enhanced. Additionally, Tsotsos and colleagues (*1995, 2008*) proposed the selective tuning model, which directly suggested that the inhibitory zone surrounding the attended item results from top-down propagation of a winner-take-all mechanism that attenuates irrelevant upstream connections iteratively from one hierarchical level down to the next. Thus, without the distractor, these irrelevant upstream connections across the visual cortical processing hierarchy could be eliminated or strongly attenuated, which, in turn, affects the inhibitory zone surrounding the attended target, in other words, resulting in the gradient rather than center-surround profile of attentional modulation.

### A wider scope of visual bottom-up attention with than without awareness

Our study provided convening evidence for a wider scope of visual bottom-up attention with than without awareness. Although this result cannot be explained by the difference in the cueing effect between visible and invisible conditions (*Figure 2D*), it could be argued that this result can be derived by that the visible relative to invisible condition has involved some degree of endogenous attention as well. Particularly, for Experiment 1 (VCCP, *Figure 1A*), compared to the invisible condition, subjects during the visible condition knew on each trial that the exogenous cue appeared randomly across the possible 9 positions and could have therefore directed endogenous attention to all, which might increase the attentional set (*Couperus and Lydic, 2019; Gibson and Kelsey, 1998*), thus yielding a wider scope of attentional modulation. Critically, it is important to note that in our study, the task required subjects to discriminate the orientation (45° or 135°) of the probe; the exogenous cue was never task-relevant. Thus, subjects did not need to direct endogenous attention to these task-irrelevant cues. More importantly, this endogenous attention (i.e., the attentional set) explanation couldn’t account for the same qualitative conclusion in our Experiment 2 (CCVP, *Figure 1D*) since the same exogenous cue was always presented at the center of 9 locations during both the visible and invisible conditions. In other words, there was the same attentional set between two conditions. If the wider scope of visual bottom-up attention with than without awareness is derived by this potential endogenous attention (i.e., the attentional set driven by the observer’s knowledge of the cue location), then we should have observed a similar scope of the two. However, our data show that this is not the case.

Our findings can be viewed as identifying an awareness-dependent scope of visual bottom-up attention. Note that this conclusion is based on a report-based paradigm in which subjects overtly push a button to report their percept. Several studies have argued that such report-based paradigms could be modulated by factors that are not directly related to the scope of attentional modulation, such as higher-level strategies, response history, experience, learning, response biases, and personality (*Yeshurun, 2019*). In addition, using a no-report paradigm, such as recording subjects’ pupillary light responses, Tkacz-Domb and Yeshurun (*2018*) revealed that the scope of attentional modulation was twofold larger than that estimated using the traditional report-based paradigm. It is important to note that subjects in our study performed exactly the same task between visible and invisible conditions (*Figure 1C and 1F*), thus the awareness-dependent scope of visual bottom-up attention evident here cannot be explained by this discrepancy between the report-based and no-report paradigms. However, the underlying neural basis of awareness-dependent scope of visual bottom-up attention could depend on whether subjects overtly report their percept.

On the one hand, several theories of conscious awareness, including the neuronal global workspace (*Dehaene and Changeux, 2011; Mashour et al., 2020*), information integration (*Koch et al., 2016; Tononi et al., 2016*), and higher-order (*Lau and Rosenthal, 2011*) theories propose that the neural activity in frontoparietal cortex is essential for conscious awareness. Similar to our study, evidence from those theories typically used the report-based paradigm in which subjects overtly report their percept, and showed that a broad frontoparietal network of areas could be activated during various tasks that contrast perceived stimuli with invisible stimuli (*Dehaene and Changeux, 2011; Mashour et al., 2020*). Thus, although speculative, it is plausible that the wider scope of bottom-up attention with than without awareness evident in our study may result from the increased activity in frontoparietal cortical areas. On the other hand, several studies have argued that such report-based paradigms do not dissociate the brain regions required for pure conscious experience from those involved in conscious access and reportability (*Koch et al., 2016; Tsuchiya et al., 2015*). Those studies, by contrast, found that posterior rather than frontoparietal cortical areas were activated when using a no-report paradigm, such as recording eye movements and pupil dilation (*Aru et al., 2012; Frässle et al., 2014*). In other words, the awareness-dependent scope of visual bottom-up attention is more likely to be mediated by posterior cortical areas when using no-report paradigm in which there is not any overt report. Consequently, further work is needed using both report-based and no-report paradigms to examine the difference in scope of visual bottom-up attention with and without awareness, as well as their distinct neural mechanisms.

### Awareness-dependent normalization framework of visual bottom-up attention

The most parsimonious account of our results is that visual bottom-up attention interacts with the normalization processes depending on awareness. Importantly, this result cannot be explained by a number of factors, such as the strength of cueing effect, post-stimulus cue, or an involvement of endogenous attention. First, both the visible and invisible cues were the same with those in Distribution Experiments and no significant difference in cueing effect was found between the two (*Figure 2D*). Second, although previous studies have suggested that the post-stimulus cue (for example, the response cue in our study) can influence not only subjects’ non-perceptual decision (*Eckstein et al., 2013*) but also the perception of stimuli presented before it (*Pestilli et al., 2011; Sergent et al., 2013*), the response cue in our study was totally randomized and uninformative about the target grating; we thus believe that our psychophysical results cannot be explained by the response cue. Finally, subjects knew before each trial that the discrimination task was to be performed on one of two gratings and could have therefore directed endogenous attention to both, and thus it is not known exactly how exogenous attention and this endogenous attention combine (*Herrmann et al., 2010*). However, subjects in our study performed exactly the same task between visible and invisible conditions, this potential combination thus couldn’t account for the observed awareness-dependent normalization processes of visual bottom-up attention.

Our data can be interpreted by a hypothesis that behavioral performance is limited by the neuronal activity with an additive, independent, and identically distributed noise, and the decision-making process with a maximum-likelihood decision rule (*Jazayeri and Movshon, 2006; Pestilli et al., 2009*). Performance accuracy *d’*, used in both previous (*Herrmann et al., 2010; Zhang et al., 2016*) and our studies here, is proportional to the signal-to-noise ratio of the underlying neuronal responses. Thus, it can parallel reflect any change in neuronal CRFs in our study. Indeed, we found that a change in the cueing effect consonant with a change in contrast gain of CRF for bottom-up attention with awareness and a change in response gain of CRF for bottom-up attention without awareness (*Figure 4A*). These awareness-dependent gain modulations of visual bottom-up attentional selection support and extend the normalization model of attention (*Boynton, 2009; Carandini and Heeger, 2012; Herrmann et al., 2010; Lee and Maunsell, 2009, 2010; Reynolds and Heeger, 2009; Reynolds et al., 1999*). This model proposes that, in the absence of attention (e.g., in the incongruent cue condition), two factors determine the firing rate of a visually responsive neuron. One is the stimulus drive (excitatory component) determined by the contrast of the stimulus placed in the receptive field of a neuron. The other is the suppressive drive (inhibitory component) determined by the summed activity of other neighboring neurons, which serves to normalize the overall spike rate of the given neuron via mutual inhibition (*Heeger, 1992; Reynolds and Heeger, 2009*). Attention (e.g., in the congruent cue condition) modulates the pattern of neural activity by altering the balance between these excitatory and inhibitory components, depending on the relative sizes of the attention field to the stimulus size, and thereby exhibiting response gain changes, contrast gain changes, and various combinations of the two. In our study, given the scope of visual bottom-up attention was significantly wider with than without awareness (*Figure 2*), for each subject, the size of the target stimuli in the spatial cueing task was manipulated as their respective mean scopes of visual bottom-up attention with and without awareness (*Figure 3A*). Thus relative to the stimulus size, the broadened attention field by visible exogenous cues led to contrast gain changes because attentional gain was applied equally to the stimulus and suppressive drives. Conversely, the narrowed attention field by invisible exogenous cues led to response gain changes because attentional gain enhanced the entire stimulus drive, but only enhanced the center of the suppressive drive. Indeed, using the classical normalization model of attention, we successfully simulated these broadened and narrowed attention fields of visible and invisible cues, respectively (*Figure 5*), further supporting an awareness-dependent normalization framework of visual bottom-up attention.

Notably, evidence from neurophysiological and brain imaging studies indicate controversies concerning the brain regions involved in visual bottom-up attention, such as subcortical structures (*Fecteau and Munoz, 2006; Shipp, 2004*), visual (*Mazer and Gallant, 2003; Zhang et al., 2012*) and frontoparietal (*Bisley and Goldberg, 2010; Corbetta and Shulman, 2002; Moore and Zirnsak, 2017; Squire et al., 2013*) cortical areas. An important factor of this controversy is the awareness, which determines whether the realized neural substrate reflects the pure bottom-up attention or not (*Chen et al., 2016; Huang et al., 2020; Zhang et al., 2012*). Intriguingly, our results are consistent with this idea by showing an awareness-dependent normalization framework of visual bottom-up attention. Although normalization as a neural computation likely occurs throughout the whole brain (*Carandini and Heeger, 2012; Schmitz and Duncan, 2018*), the observed neural correlates of its interaction with visual bottom-up attention could also depend on awareness, and further studies will shed light on this issue using neurophysiological or brain imaging techniques.

## Conclusions

In sum, we conclude that manipulating subjects’ awareness can modulate the scope of visual bottom-up attentional modulation, which, in turn, affects its normalization processes. Our study provides, to the best of our knowledge, the first experimental evidence supporting an awareness-dependent normalization framework of visual bottom-up attention, thereby furthering our understanding of the neural computations underlying visual attention, the relationship between attention and awareness, and how they interactively shape our experience of the world.

## Methods and Materials

### Subjects

A total of 16 human subjects (8 male, 19–26 years old) were involved in the study. All of them participated in Distribution Experiments, fourteen of them repeated the Distribution Experiments with decreased luminance of visible cues, and the following Normalization Experiments. They were naïve to the purpose of the study. They were right-handed, reported normal or corrected-to-normal vision, and had no known neurological or visual disorders. They gave written, informed consent, and our procedures and protocols were approved by the human subjects review committee of School of Psychology at South China Normal University.

### Apparatus

Visual stimuli were displayed on an IIYAMA color graphic monitor (model: HM204DT; refresh rate: 60 Hz; resolution: 1,280 × 1,024; size: 22 inches) at a viewing distance of 57 cm. Subjects’ head position was stabilized using a chin rest. A white fixation cross was always present at the center of the monitor.

### Distribution Experiments

#### Stimuli

As illustrated in *Figure 1*, each texture stimulus contained 18 positions (the possible locations of the exogenous cue and probe) settled at an iso-eccentric distance from fixation (8.27° of visual angle); a half of them were located in the left visual field and the other half was located in the right visual field. The center-to-center distance between two neighboring positions was 1.35°. The exogenous cue was a low-luminance ring (8.9 cd/m^2^; inner dimeter: 0.909°; outer dimeter: 0.961°) while the probe was a rectangle of 0.104° × 0.831° in visual angle and was orientated at 45° or 135° away from the vertical. Low- and high-contrast masks, which had the same grid as the texture stimulus, rendered the exogenous cue visible or invisible (confirmed by a two-alternative forced choice, 2AFC) to subjects, respectively. Each mask ring contained two pairs of orthogonal circular arcs, one pair was white (19.3 and 79.8 cd/m^2^ for Low- and high-contrast masks, respectively) and the other pair was black (11.3 and 0.01 cd/m^2^ for Low- and high-contrast masks, respectively). The ring in the mask had the same size as the exogenous cue in the texture stimulus (*Figure 1B and 1E*).

#### Procedure

The Distribution Experiment consisted of 3 experiments. In each visual field, the probe position was constant and the exogenous cue position varied in Experiment 1 (i.e., the varied cue with constant probe, VCCP, *Figure 1A*), whereas Experiment 2 was a converse situation (i.e., the constant cue with varied probe, CCVP, *Figure 1D*). In both Experiments 1 and 2, there were five possible distances between the exogenous cue and probe, ranging from D0 (cue and probe at the same location) through D4 (cue and probe four items away from each other). Subjects participated in Experiments 1 and 2 on two different days, and the order of the two experiments was counterbalanced across subjects. Experiment 3 checked the effectiveness of the awareness manipulation in both Experiments 1 and 2, and was always before them. In both Experiments 1 and 2, each trial began with the fixation. A cue frame with (the cue condition) or without (the non-cue condition) exogenous cue was presented for 50-ms, followed by a 100-ms mask (low- and high-contrast in visible and invisible conditions, respectively, confirmed by Experiment 3, *Figure S1*) and another 50-ms fixation interval. Then a probe line, orientating at 45° or 135° away from the vertical, was presented for 50-ms. Subjects were asked to press one of two buttons as rapidly and correctly as possible to indicate the orientation of the probe (45° or 135°). The cueing effect for each distance (D0 to D4) was quantified as the difference between the reaction time of the probe task performance in the non-cue condition and that in the cue condition.

Differently, Experiment 1 consisted of 16 blocks of 96 trials, 48 for the left visual field and 48 for the right visual field. In each block and each visual field, an exogenous cue was equiprobably and randomly presented at one of the 9 positions in 40 trials (the cue condition) and was absent in the remaining 8 trials (the non-cue condition). The probe was always presented at the center of 9 positions (i.e., the VCCP, *Figure 1A*). Experiment 2 consisted of 16 blocks of 80 trials, 40 for the left visual field and 40 for the right visual field. In each block and each visual field, an exogenous cue always appeared (the cue condition) or was absent (the non-cue condition) in the center of 9 positions with equal probability; the probe appeared equiprobably and randomly across the possible 9 positions (i.e., the CCVP, *Figure 1D*).

The stimuli and procedure in the 2AFC experiment (i.e., Experiment 3) were the same as those in Experiments 1 and 2, except that no probe was presented (*Figure S1*). Experiment 3 checked the effectiveness of the awareness manipulation in Experiments 1 and 2, and was always before them. In Experiment 3, all subjects underwent a 2AFC task to determine whether the masked cue was visible or invisible in a criterion-free way. After the presentation of a masked cue frame, subjects were asked to indicate which side (upper left or upper right) from the fixation they thought the cue appeared. Their performances were significantly higher or not statistically different from chance for all possible distances (D0 to D4), providing an objective confirmation that the cue was indeed visible or invisible to subjects, respectively.

#### Model fitting and comparison

For each subject and each condition (visible and invisible), we fitted a monotonic model and two non-monotonic models to the averaged cueing effect. The monotonic model was implemented as the Gaussian function, and the two non-monotonic models were implemented as the Mexican Hat (i.e., a negative second derivative of a Gaussian function) and Polynomial functions (*Fang and Liu, 2019; Fang et al., 2019; Finke et al., 2008*):

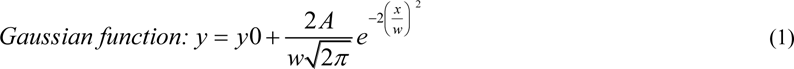

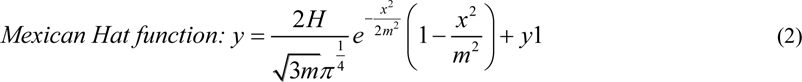

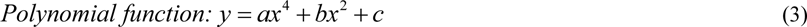

where *y* is the cueing effect, *x* is the distance between the cue and probe (i.e., D0 to D4); *w*, *A*, and *y0* are the three parameters controlling the shape of the Gaussian function; *m*, *H*, and *y1* are three parameters controlling the shape of the Mexican Hat function; *a*, *b*, and *c* are the three free parameters controlling the shape of the Polynomial function (note that we used a fourth-order polynomial without the odd-power terms for the cueing effect since the symmetric shape). To compare these three models to our data, we first computed the Akaike information criterion (AIC, *Akaike, 1973*) and Bayesian information criterion (BIC, *Schwarz, 1978*), with the assumption of a normal error distribution:

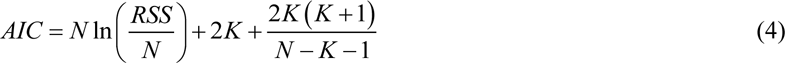

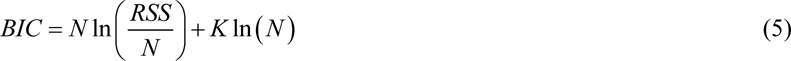

where *N* is the number of observations, *K* is the number of free parameters, and *RSS* is residual sum of squares (*Raftery, 1999*). Then, we further calculated the Likelihood ratio (LR) and Bayes factor (BF) of the monotonic model (Gaussian) over non-monotonic models (Mexican Hat and Polynomial) based on AIC (*Burnham and Anderson, 2002*) and BIC (*Wagenmakers, 2007*) approximation, respectively:

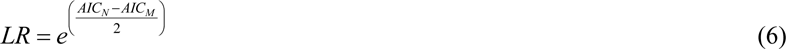

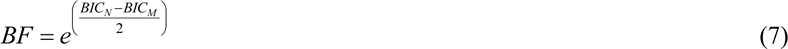

where *AIC_M_* and *BIC_M_* are for the monotonic (Gaussian) model, *AIC_N_* and *BIC_N_* are for non-monotonic (Mexican Hat and Polynomial) models. The results indicated that, during each condition (visible and invisible), the monotonic (Gaussian) model was strongly favored over the non-monotonic models (Mexican Hat and Polynomial) (*Figure 2*). Thus, to quantitatively examine the scope of attentional modulation, we fit the averaged cueing effects from D0 to D4 with a Gaussian function and used the FWHM (full width at half maximum) bandwidth of the Gaussian to quantify their scopes:

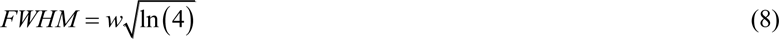

where *w* is the fitted width of the Gaussian function.

### Normalization Experiments

#### Stimuli

As illustrated in *Figure 3C*, the exogenous cue of Normalization Experiments was the same as those in the Distribution Experiment 2, i.e., the exogenous cue always appeared in the center of 9 positions in left or right hemifield at 8.27° eccentricity. The probe was a pair of gratings (spatial frequency: 1.7 cycles/°; phase: random) that were presented at the exogenous cue’s locations in the left and right hemifields. The gratings were presented at five possible contrasts: 0.02, 0.08, 0.15, 0.40, and 0.70. For each subject, the diameter of grating was manipulated as their respective mean FWHM bandwidth for the Gaussian of bottom-up attention with and without awareness in Distribution Experiments (*Figure 3A*):

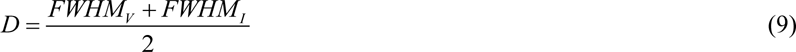

where ***D*** is the diameter of grating, *FWHM_V_* and *FWHM_I_* are the fitted FWHM bandwidth of the Gaussian model in the visible and invisible conditions during Distribution Experiments (Experiments 1 & 2), respectively.

### Procedure

The Normalization Experiment consisted of 2 experiments. Each trial began with central fixation. The exogenous cue, a low-luminance ring, randomly appeared at the center of 9 positions in left or right hemifield with equal probability, followed by a 100-ms mask (low- and high-contrast for visible and invisible conditions, respectively) and another 50-ms fixation interval. Then, a pair of gratings (with identical contrasts) was presented for 33 ms in the left and right hemifields, one of which was the target. Subjects were asked to press one of two buttons to indicate the orientation of the target grating (leftward or rightward tilted) and received auditory feedback if their response was incorrect. The target grating was indicated by a peripheral 100 ms response cue (0.4° black circular arc) above one of the grating locations, but not at the grating location to avoid masking. A congruent cue was defined as a match between the exogenous cue location and response cue location (half the trials); an incongruent cue was defined as a mismatch (half the trials) (*Figure 3C*). Subjects were explicitly told that the exogenous cue was randomized and uninformative about the target location. The Normalization Experiment consisted of two sessions (visible and invisible), with the two sessions occurring on different days; the order of the two sessions was counterbalanced across subjects. Each session consisted of 64 blocks; each block had 80 trials, from randomly interleaving 16 trials from each of the five contrasts. Contrast varied from trial to trial in randomly shuffled order, and stimuli were presented briefly (i.e., 33 ms) to avoid any possible dependence of attentional state on stimulus contrast. The attentional effect for each grating contrast was quantified as the difference between the performance accuracy (*d’*) in the congruent and incongruent cue conditions.

### Psychophysical data analysis

To quantitatively examine the pattern of gain (either contrast or response gain) separately for bottom-up attention with and without awareness, for each subject, performance—i.e., *d’* = z (hit rate) - z (false alarm rate)—was assessed across experimental blocks for each contrast and each trial condition (congruent and incongruent). A rightward response to a rightward stimulus tilt was (arbitrarily) considered to be a hit, and a rightward response to a leftward stimulus was considered to be a false alarm. For each subject, the mean *d’* CRFs obtained for congruent and incongruent trials were fit with the standard Naka–Rushton equation (*Naka and Rushton, 1966*):

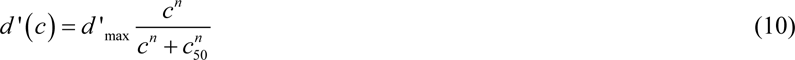

where *d’* is performance as a function of contrast (c), *d’* _max_ determines the asymptotic performance at high contrasts, *c_50_* is the contrast corresponding to half the asymptotic performance, and *n* is an exponent that determines the slope of the CRFs. The two parameters *d’* _max_ and *c_50_* determined response gain and contrast gain, respectively. We estimated these two parameters for each condition while *n* (slope) was fixed at 2 according to previous studies (*Carandini and Heeger, 2012; Herrmann et al., 2010; Reynolds and Heeger, 2009; Zhang et al., 2016*).

### Model simulations

The normalization model of attention (*Reynolds and Heeger, 2009*) computes the response of an arbitrary single neuron to a given set of stimuli as:

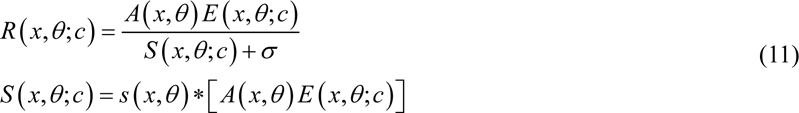

where *R(x,θ;c)* is the response of a neuron with its receptive field centered at *x* and its orientation tuning centered at *θ*, receiving stimulus input with contrast *c*, *A(x,θ)* is the attention field, *E(x,θ;c)* is the stimulus drive of the population of neurons evoked by contrast *c*, *σ* is the normalization constant*, S(x,θ;c)* is the effect of the normalizing pool and represents the excitatory drive convolved by the suppressive surround, *s(x,θ)* is the suppressive field, and *\** is convolution. Applying the attention field in the model can yield either a change in response gain, a change in contrast gain, or a combination of the two, depending on the stimulus size and the extent of the attention field relative to the sizes of the stimulation and suppressive fields. Relative to the stimulus size, the broadened attention field led to contrast gain changes since attentional gain (*λ*) was applied equally to the stimulus (*αc*, *α* is the constant gain of the neuron receiving it’s preferred input with contrast *c*) and suppressive drives (*S*). The responses of a model neuron can be approximated as:

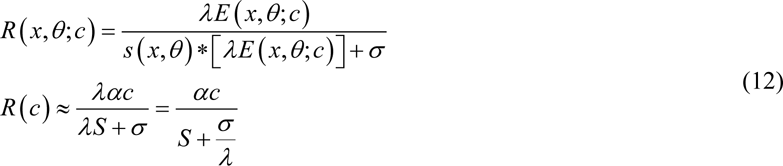

Conversely, the narrowed attention field led to response gain changes since attentional gain (*λ*) enhanced the entire stimulus drive (*αc*), but its impact on the denominator *S* + *σ* is much minimal. In this case:

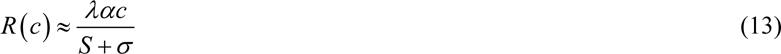

Our results supported these predictions of the normalization model of attention by showing that manipulating subjects’ awareness could modulate the field of visual bottom-up attention, which, in turn, affected its normalization processes (see *Figure 4*). To further confirm this awareness-dependent normalization framework of visual bottom-up attention, we simulated our empirical data using custom Matlab scripts based on the code of Reynolds and Heeger (*2009*) with 4 free parameters: the gain of attention [*A(x,θ)*], separately optimized for visible and invisible cue conditions, the normalization constant *σ*, and a scaling parameter to linearly scale simulated values to *d’*. Given the simulated attention fields [*A*(*x*,*θ*)] are in arbitrary units; only the relative values are meaningful (*Reynolds and Heeger, 2009*), in both the visible and invisible conditions, we thus calculated the correlation coefficients between the simulated attention field and experimental attention fields (i.e., the FWHM) across individual subjects (see *Figure 5*).

## Acknowledgements

We thank David Heeger and Ruyuan Zhang for valuable comments. This work was supported by the National Outstanding Youth Science Fund Project of National Natural Science Foundation of China (Project: 32022032), the National Natural Science Foundation of China (General Program: 31871135), and the Key Realm R&D Program of Guangzhou (202007030005).

## Additional information

### Funding

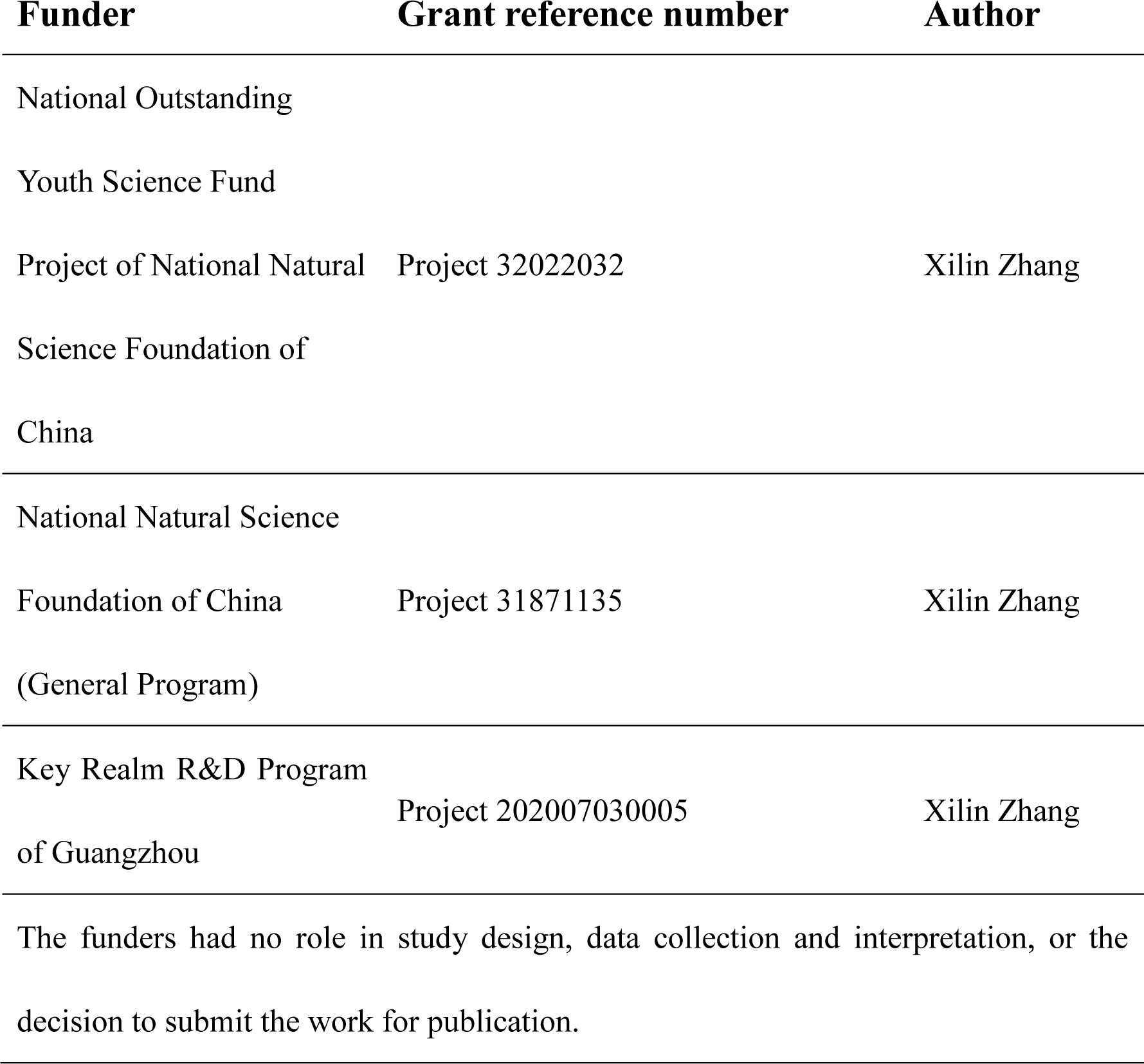

### Author contributions

Shiyu Wang, Conceptualization, Formal analysis, Investigation, Methodology, Writing – original draft, Writing - review and editing; Ling Huang, Qinglin Chen, Jingyi Wang, Siting Xu, Formal analysis, Investigation, Methodology; Xilin Zhang, Conceptualization, Formal analysis, Supervision, Funding acquisition, Investigation, Methodology, Writing - original draft, Writing - review and editing

### Ethics

Human subjects: They study was approved by the human subjects review committee of School of Psychology at South China Normal University.

## Additional files

### Data availability

The data and custom-built MATLAB scripts of this study are available at Open Science Framework: https://osf.io/gqzvm/.

## Supplementary data

**Figure S1.**
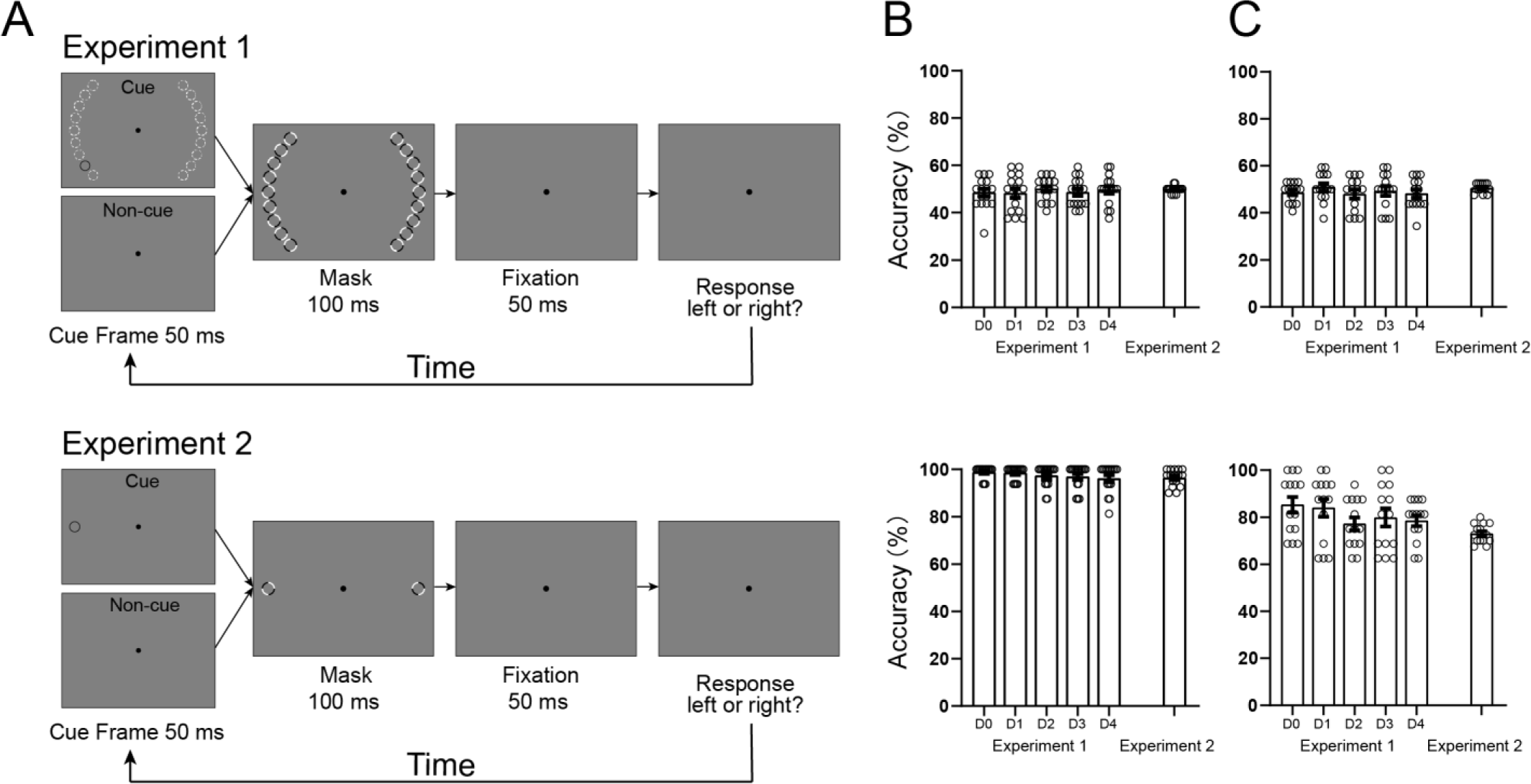
Protocol and Results of the Two-alternative Forced-choice Test. **(A)** The stimuli and procedure in the two-alternative forced-choice (2AFC) experiment (i.e., Experiment 3) were the similar to those in Experiments 1 (top) and 2 (bottom), except that no probe was presented. Experiment 3 checked the effectiveness of the awareness manipulation in Experiments 1 and 2, and was always before them. In Experiment 3, all subjects underwent a 2AFC task to determine whether the masked cue was visible or invisible in a criterion-free way. After the presentation of a masked cue frame, subjects were asked to indicate which side (left or right) from the fixation they thought the exogenous cue appeared. Their performances were significantly higher or not statistically different from chance, providing an objective confirmation that the exogenous cue was indeed visible or invisible to subjects, respectively. **(B)** For the invisible condition (top), subjects reported that they were unaware of the exogenous cue and could not detect which visual filed contained it. Their performances were not statistically different from chance [mean percent correct ± standard error of the mean, Experiment 1 (i.e., varied cue), D0: 47.656 ± 1.651%, D1: 48.242 ± 1.996%, D2: 49.609 ± 1.303%, D3: 45.508 ± 2.096%, D4: 49.609 ± 1.533%, all t_15_ < 0.968, *p* > 0.348, 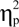 < 0.500; Experiment 2 (i.e., constant cue): 49.503 ± 0.411%, t_15_ = 0.436, *p* = 0.669, 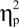 = 0.225]; for the visible condition, by contrast, their performance was significantly higher than chance (Experiment 1, D0: 98.828 ± 0.630%, D1: 98.438 ± 0.699%, D2: 97.266 ± 1.137%, D3: 96.875 ± 1.276%, D4: 96.094 ± 1.496%, all t_15_ > 30.812, *p* < 0.001, 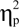 > 15.911; Experiment 2: 99.503 ± 0.411%, t_15_ = 52.557, *p* < 0.001, 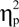 = 27.140). Furthermore, for the Experiment 1, subjects’ performances were submitted to a repeated-measures ANOVA with awareness (visible and invisible) and distance (D0 to D4) as within-subjects factors. The main effect of distance (F_4, 60_ = 0.215, *p* = 0.929, 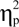 = 0.014) and the interaction between the two factors (F4, 60 = 0.942, *p* = 0.446, 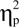 = 0.059) were not significant, but the main effect of awareness was significant (F_1, 15_ = 1873.86, *p* < 0.001, 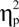 = 0.992). These results indicate that our awareness manipulation was effective for both the visible and invisible conditions, and there was no significant difference in subject performance among five distances. Error bars denote 1 SEM calculated across subjects and open dots denote the data from each subject. **(C)** To manipulate the cueing effect between visible and invisible conditions, we decreased the luminance of the cue in visible condition. Fourteen of our 16 subjects repeated the same 2AFC test and the results showed that subjects’ performances were not statistically different from chance in the invisible condition [Experiment 1 (i.e., varied cue), D0: 48.661 ± 1.121%, D1: 50.893 ± 1.652%, D2: 47.991 ± 1.927%, D3: 49.330 ± 2.050%, D4: 48.214 ± 1.725%, all t_13_ < 1.194, *p* > 0.254, 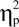 < 0.662; Experiment 2 (i.e., constant cue): 50.536 ± 0.536%, t_13_ = 1.000, *p* = 0.336, 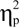 = 0.555], but were significantly higher than chance in the visible condition (Experiment 1, D0: 85.268 ± 3.315%, D1: 83.929 ± 3.747%, D2: 77.232 ± 2.749%, D3: 79.911 ± 3.774%, D4: 78.571 ± 2.336%, all t_13_ > 7.926, *p* < 0.001, 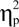 > 4.397; Experiment 2: 73.036 ± 1.086%, t_13_ = 21.207, *p* < 0.001, 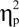 = 11.764). Similarly, for the Experiment 1, subjects’ performances were also submitted to a repeated-measures ANOVA with awareness (visible and invisible) and distance (D0 to D4) as within-subjects factors. The main effect of distance (F_4, 52_ = 1.892, *p* = 0.126, 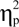 = 0.127) and the interaction between the two factors (F4, 52 = 0.923, *p* = 0.458, 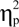 = 0.066) were not significant, but the main effect of awareness was significant (F_1, 13_ = 171.254, *p* < 0.001, 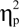 = 0.929). These results further conform that our awareness manipulation was effective for both the visible and invisible conditions, and there was no significant difference in subject performance among five distances. Error bars denote 1 SEM calculated across subjects and open dots denote the data from each subject.

**Figure S2.**
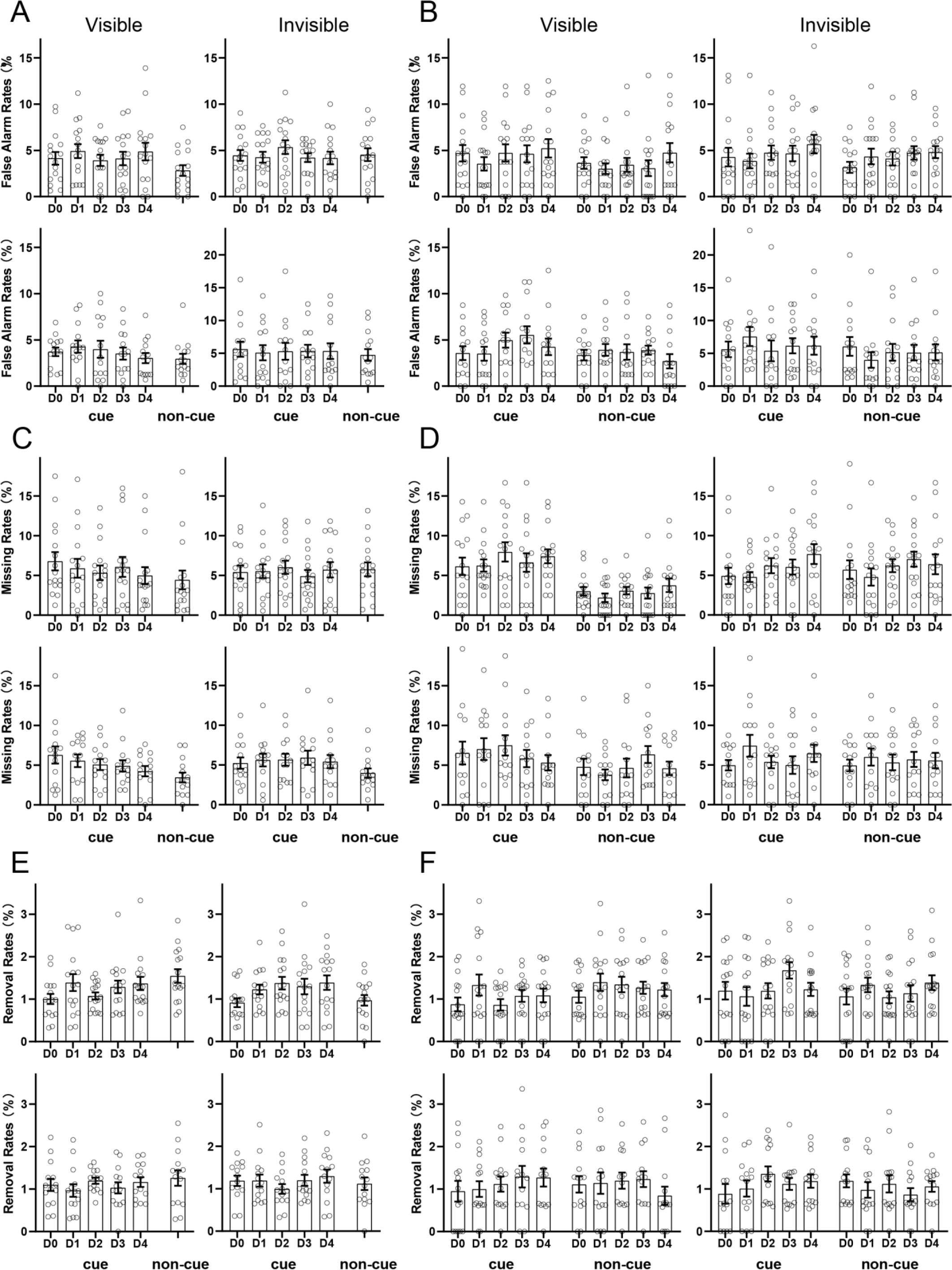
False alarm, Miss, and Removal Rates in Distribution Experiments. False alarm rates of the cue and non-cue conditions for each distance (D0 to D4) during visible (left) and invisible (right) conditions in Experiments 1 **(A)** and 2 **(B)**. Note that, in our study, subjects were asked to press one of two buttons as rapidly and correctly as possible to indicate the orientation of the line probe (45° or 135°). Thus, for each condition, a rightward response to a 45° line was (arbitrarily) considered to be a hit, a rightward response to a 135° line was considered to be a false alarm, and a leftward response to a 45° line was considered to be a miss. Error bars denote 1 SEM calculated across subjects and open dots denote the data from each subject. Miss rates of the cue and non-cue conditions for each distance (D0 to D4) during visible (left) and invisible (right) conditions in Experiments 1 **(C)** and 2 **(D)**. Error bars denote 1 SEM calculated across subjects and open dots denote the data from each subject. Removal rates (i.e., correct reaction times shorter than 200 ms and beyond three standard deviations from the mean reaction time in each condition were removed) of the cue and non-cue conditions for each distance (D0 to D4) during visible (left) and invisible (right) conditions in Experiments 1 **(E)** and 2 **(F)**. Error bars denote 1 SEM calculated across subjects and open dots denote the data from each subject. There was no significant difference in false alarm rate, miss rate, or removal rate across conditions (all *P* > 0.05).

**Figure S3.**
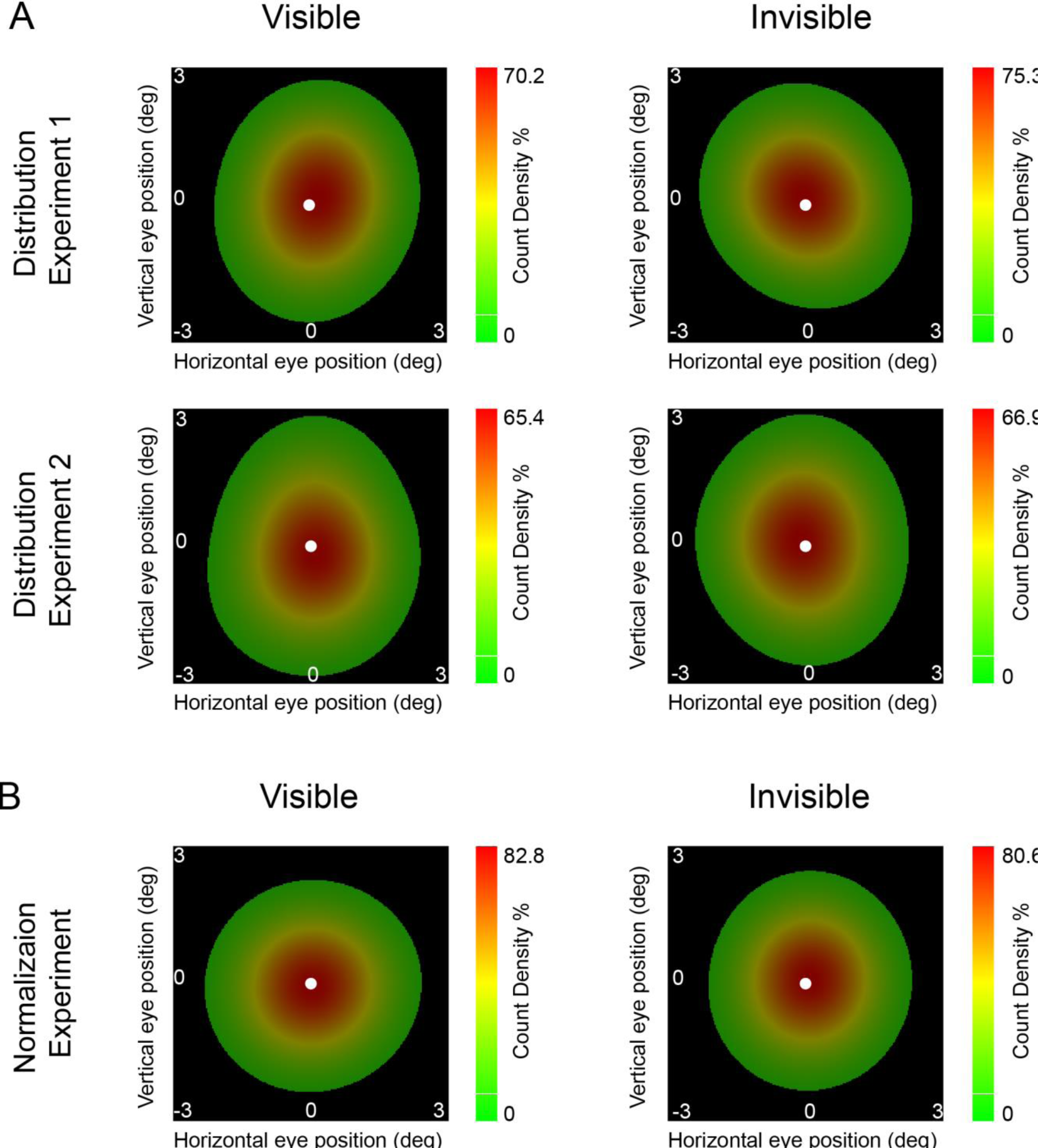
Eye Movement Data in Distribution and Normalization Experiments. Horizontal and vertical eye positions after removing blinks and artifacts of the visible (left) and invisible (right) conditions in Distribution **(A)** and Normalization **(B)** Experiments. Subjects‘ eye movements were small (< 3°) and not systematically different between the visible and invisible conditions (all *p* > 0.05).

**Figure S4.**
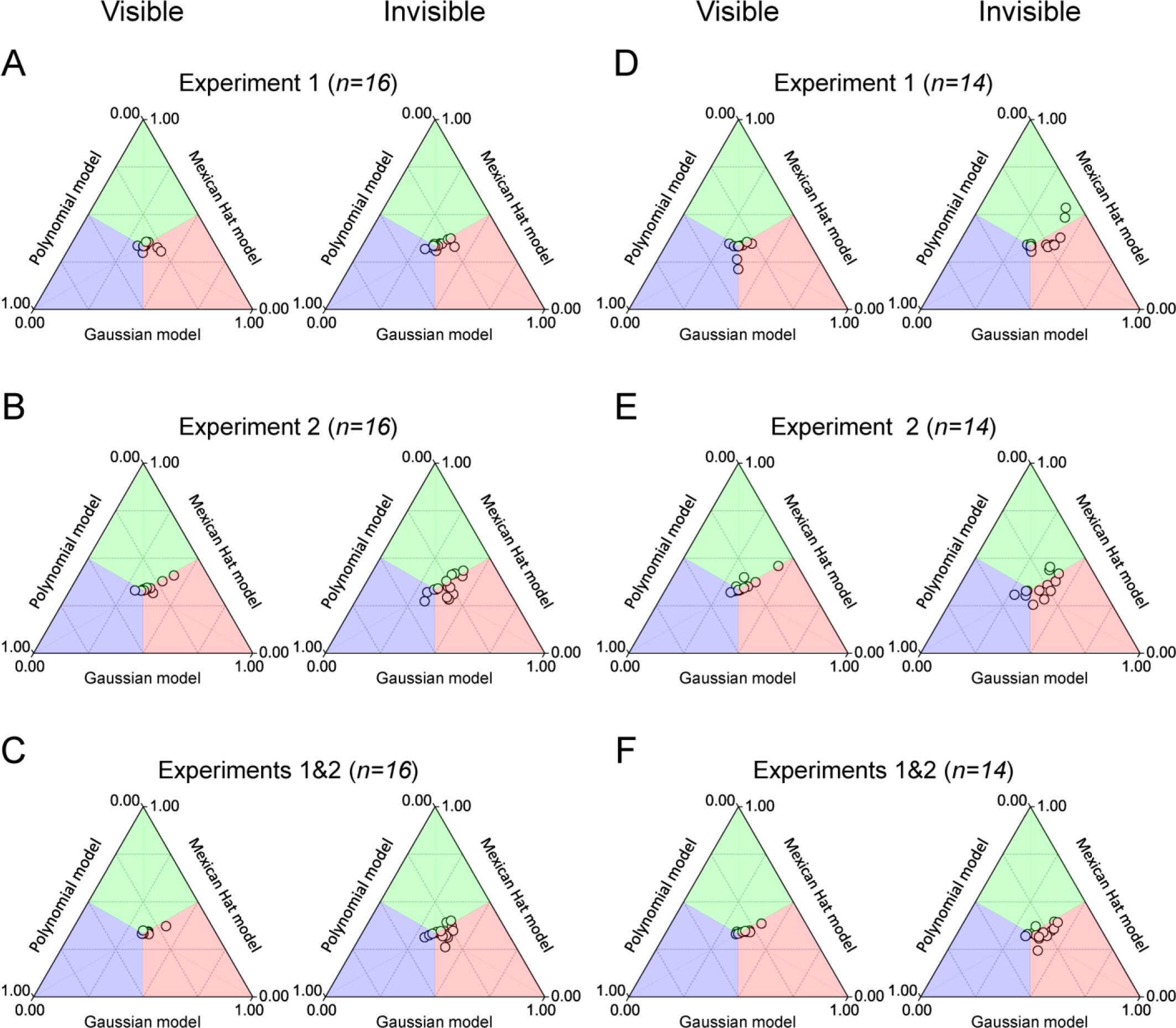
Normalized R^2^ of Individual Subjects in Distribution Experiments. In Experiment 1 (**A** and **D**), Experiment 2 (**B** and **E**), and Experiments 1 & 2 (**C** and **F**), to directly compare the fitted R^2^ among Gaussian, Mexican Hat, and Polynomial models, we normalized the R^2^ of each subject to 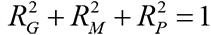, where 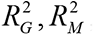 and 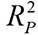 was the fitted R^2^ using the Gaussian, Mexican Hat, and Polynomial functions, respectively. During each condition (left: visible; right: invisible) and each experiment, most of the dots are located in the red zone, demonstrating that the Gaussian model was favored over both the Mexican Hat and Polynomial models.

**Figure S5.**
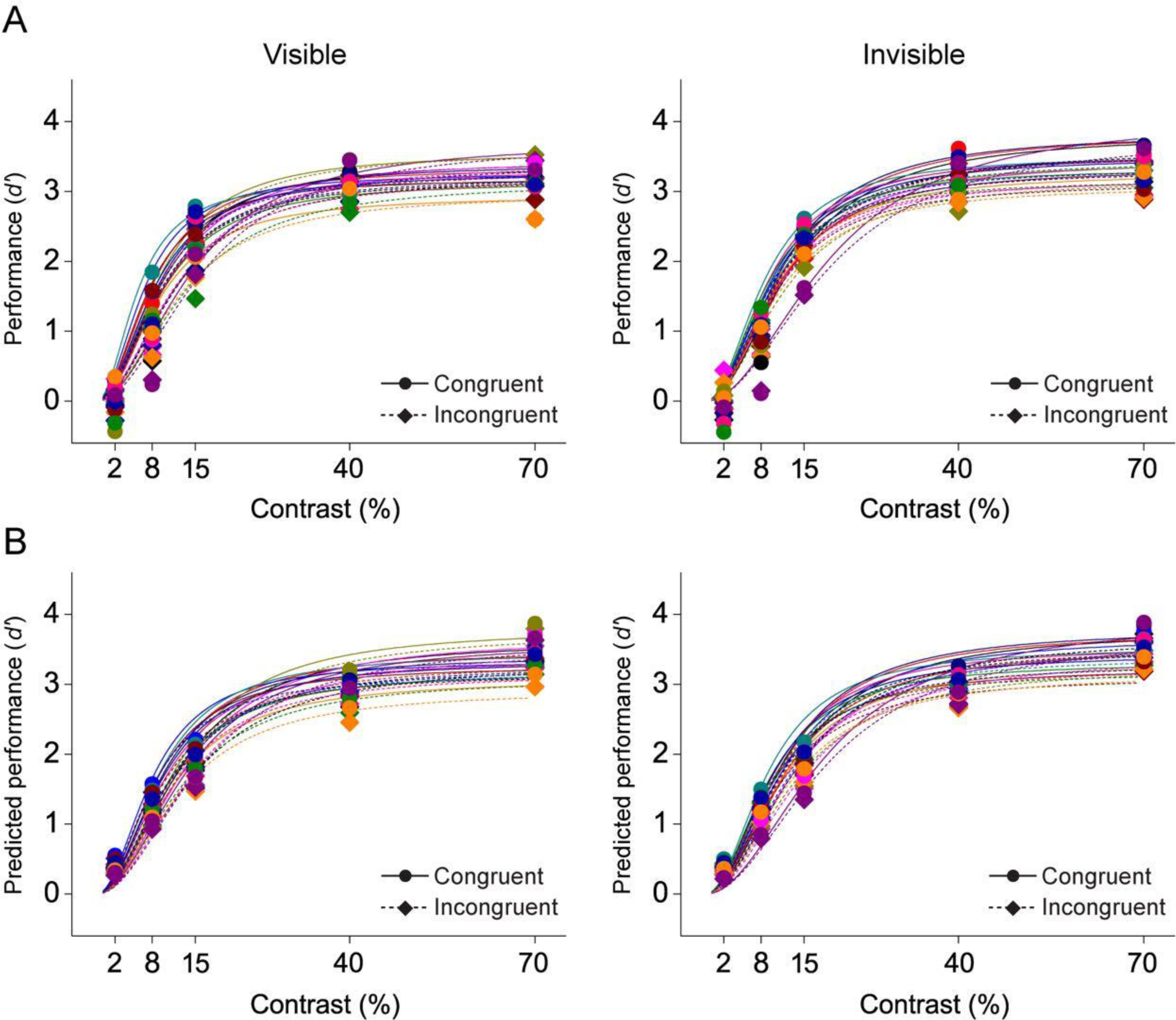
Contrast Response Functions of Individual Subjects in Normalization Experiments. Data points are experimental **(A)** and predicted **(B)** performances (*d’*) of individual subjects for awareness (visible and invisible), cue validity (congruent and incongruent), and stimulus contrasts. The individual contrast response function was fitted using the standard Naka–Rushton equation **(A)** and the original normalization model of attention with 4 free parameters **(B)**, respectively.

## Notes

### Competing Interest Statement

The authors have declared no competing interest.

